# Ligand Binding Kinetics, Thermodynamics, and Gating of Insect Odorant Receptor

**DOI:** 10.64898/2026.01.19.700284

**Authors:** Tiefeng Song, Minying Low, Huixia Shou, Haohao Fu, Yong Wang

## Abstract

Insect odorant receptors (ORs) are unique ligand-gated ion channels specialized in detecting volatile molecules. The ancestral M. hrabei odorant receptor MhOR5 functions as a homomer, operating independently of the highly conserved odorant receptor co-receptor (Orco). While MhOR5 exhibits a broad ligand-binding capacity, the molecular underpinnings of its ligand specificity and gating dynamics remain poorly understood. To resolve this knowledge gap, we applied an integrated molecular dynamics enhanced sampling approach, combining collective variable machine learning, alchemical free energy calculations, and kinetic calculations, to dissect the ligand recognition and gating mechanisms of MhOR5 in complex with two key ligands: eugenol (EOL) and N,N-diethyl-m-toluamide (DEET). Our findings reveal striking differences in the binding modes of EOL and DEET, identify dual ligand binding/release pathways (solvent-facing and membrane-facing), and pinpoint two critical molecular determinants: W158 as a central gating residue regulating ligand dissociation, and a proline-disrupted S4 helix that modulates ligand specificity via water-mediated hydrogen bonding with DEET. Thermodynamic and kinetic calculations further demonstrate that DEET binds MhOR5 substantially more stably (-11.2 kcal/mol vs. -8.4 kcal/mol for EOL) and associates marginally faster, while EOL dissociates two to three orders of magnitude more rapidly. Collectively, these findings elucidate the atomic-level mechanisms of MhOR5’s ligand recognition and gating, providing a foundational framework for understanding insect OR function.

## Introduction

Organisms require the ability to sense volatile small molecules and respond accordingly. ^1^ The vast chemical diversity of small molecules in natural environments presents a significant challenge for the chemosensory system. A common strategy employed by organisms to overcome this challenge involves the co-encoding of chemical sensation by a large family of chemoreceptors. This allows robust chemoreceptors to navigate the chemical landscape effectively. However, chemical receptors must balance broad-spectrum and specific affinity to detect odorants while remaining insensitive to others. Understanding these chemoreceptors at the molecular level could inform the design of improved attractants and repellents, potentially targeting specific pest species while minimizing impact on non-target organisms. Three primary classes of chemoreceptors have been identified: odorant receptors (ORs), gustatory receptors (GRs), and ionotropic receptors (IRs). ^2,3^ ORs are unique heteromeric ligand-gated ion channels situated in the plasma membrane of insect odorant receptor neurons (ORNs).^4,5^ While GRs are homomeric ligand-gated ion channels that are closely related to ORs,^6–8^ IRs resemble ionotropic glutamate receptors.^9^

In mammals, ORs belong to the class of G-protein-coupled receptors (GPCRs).^10,11^ In contrast, insects employ a distinct class of ligand-gated ion channels, which form a tetramer. Each subunit features seven transmembrane helices. Notably, the orientation of the seven transmembrane helices of OR is opposite to that of GPCR, with the short intracellular N-terminus and extracellular C-terminus. In general, insect ORs are heterotetramers, consisting of two types of subunits.^12^ One is a highly conserved co-receptor, Orco, which is essential for OR assembly and function.^13,14^ The other is a highly divergent OR, designated ORx, which contains the binding site for chemical signaling molecules and exhibits high chemical sensitivity within the heteromeric complex.^15^ The four monomers assemble in a square configuration, with an ion channel at the center formed by the C-terminal S7 helices of each monomer.

In addition to the extensive physiological and biochemical data accumulated on ORs, advancements in structural biology have significantly deepened our understanding of their atomic details. Notably, structural determinations using cryo-EM have been achieved for several insect ORs: the jumping bristletail *M. hrabei* OR5 homotetramer (MhOR5),^16^ the parasitic fig wasp *A. bakeri* Orco homotetramer,^17^ and a series of OR-Orco heterocomplexes recently resolved, including the pea aphid *A. pisum* ApisOR5 and ApisOR22,^18,19^ the disease-vector mosquito *A. aegypti* AaegOR10, *A. gambiae* AgamOR28,^20^ *D. melanogaster* DmOR67d and *D. bipectinata* DbOR67d.^21^ Collectively, these structures have established a conserved 1ORx:3Orco heterotetrameric assembly as the canonical architecture of insect odorant receptor complexes. These structures provide atomic insights such as the binding pocket, protein assembly interface, and ion channel of insect ORx and Orco complexes. Moreover, recent structural resolutions of a few GRs, such as *B. mori* BmGr9, and *D. melanogaster* DmGr43a and DmGr64a,^6–8^ underline the striking structural and functional similarities between GRs and ORs, suggesting closely related underlying ligand recognition mechanisms.

MhORs are considered the most ancestral members of the insect OR family, predating the emergence of Orco, and are capable of functioning independently of Orco. ^22^ MhOR5 demonstrates activity against tens of compounds with diverse chemical characteristics, spanning from eugenol to propyl acetate.^16^ Despite its homotetrameric OR topology and lack of Orco, the ligand-binding mechanism observed in MhOR5 is valuable for understanding other insect ORs. Del Mármol et al. elucidated the cryo-EM structures of MhOR5 in three states: unbound (MhOR5*_apo_*), bound to eugenol (MhOR5-EOL), and bound to N,N-diethyl- 3-methylbenzamide (MhOR5-DEET) (Fig. 1a). These structures revealed that ligand binding triggers the opening of the ion channel at the tetramer’s center, allowing ion flux into the cell and initiating an action potential. However, the binding pockets in the cryo-EM structures are deeply buried within the protein, with no apparent tunnels connecting to the protein surface (Fig. 1b). Consequently, the pathways for volatile compounds to enter or exit the binding site, as well as the associated thermodynamics, kinetics, and rate-determining steps, remain largely undisclosed.

**Figure 1:**
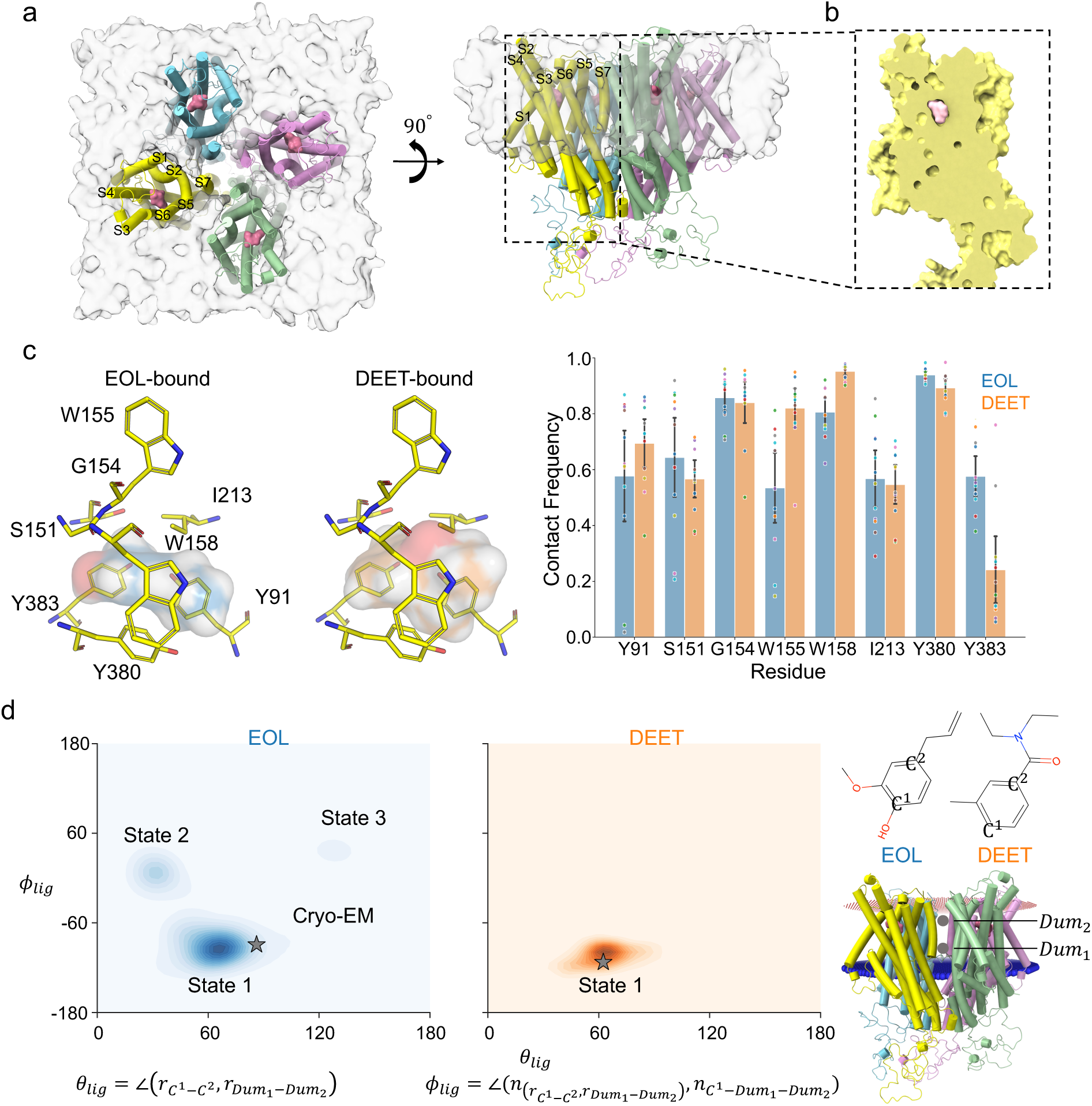
Binding pocket and binding modes of EOL and DEET. a) Surface and cartoon representation of the MhOR5-ligand complex structure, highlighting the binding pocket (left, top-down view; middle, side view after 90° rotation). b) The inset shows a close-up of the buried hydrophobic binding pocket with bound DEET. c) Close-up of the EOL-bound and DEET-bound binding pockets, with key interacting residues (yellow sticks) and ligand (pink spheres) shown; transparent surfaces indicate the ligand-binding interface. The bar graph (right) quantifies the contact frequency of each residue with EOL and DEET observed in MD simulations, demonstrating differential interaction patterns. The twelve independent trajectories are marked with scatter points of distinct colors, and the 95% CI for each trajectory is calculated to serve as error bars. d) Two-dimensional probability distributions of *θ_lig_* and *ϕ_lig_* for EOL and DEET. The ligand poses in cryo-EM data are marked as gray stars.

To gain a comprehensive understanding of ligand binding and unbinding in MhOR5, we established an integrated computational workflow spanning structural, kinetic, and thermodynamic analyses.^23^ We first used unbiased MD simulations to characterize the binding modes of EOL and DEET and assess their generality across three physicochemically related aromatic ligands. We then performed high-concentration spontaneous binding simulations to identify ligand association pathways and estimate association rates. Ligand dissociation was subsequently investigated using Random Acceleration Molecular Dynamics (RAMD),^24^ Adiabatic-bias Molecular Dynamics (ABMD), ^25^ and On-the-fly Probability Enhanced Sampling Flooding (OPES-Flooding), ^26^ guided by the AI-driven collective variable learning algorithm Koopman-reweighted time-lagged independent component analysis (KTICA),^27^ to resolve unbinding pathways, identify the W158 gating mechanism, and estimate dissociation rates. Free energy perturbation (FEP) calculations^28^ provided the final thermodynamic assessment by quantifying absolute binding free energies. Together, this workflow connected binding modes, association and dissociation pathways, gating dynamics, kinetics, and thermodynamics within a unified framework.

## Results

### Unbiased MD Reveals Subtle Binding Modes of Odorants

To elucidate the binding modes of both EOL and DEET, three all-atom unbiased MD simulations were performed, each lasting one microsecond. Since MhOR5 functions as a homotetramer composed of four identical subunits, the total effective simulation time reached 12 *µ*s. Error estimation and autocorrelation analyses confirmed that the simulations reached satisfactory equilibration (Figs. S1–S7). The simulations showed that both EOL and DEET remained fully enclosed and relatively stable within the binding pocket throughout the simulations (Fig. S8). Both ligands formed extensive van der Waals interactions with MhOR5 in a largely similar fashion, although notable differences were evident around W155, W158 and Y383 (Fig. 1c). Notably, EOL sometimes formed hydrogen bonds with S151, whereas DEET exhibited no direct hydrogen bonding interactions (Fig. S9).

To investigate the orientation dynamics of EOL and DEET, we projected the trajectories onto two CVs: the angle *θ_lig_* between the principal axis of each ligand and the membrane normal, and the dihedral *ϕ_lig_* (see Method for definitions). The resulting probability distributions revealed a primary conformational state (State 1) for both ligands which aligns with the orientation observed in the cryo-EM structures (Figs. 1d and S10). Beyond this common state, EOL sampled a broader conformational landscape, exhibiting two additional lower-populated states (State 2 and 3, Fig. 1d).

Further structural analysis indicates that the dominant State 1 corresponds to an “upward-facing” ligand conformation for both odorants (Fig. S10). In this pose, both ligands primarily interact with W155 and W158, with weaker interactions with Y383. For EOL, the *∼*30*^◦^* upward-facing conformation (State 2) is more vertical, allowing van der Waals interations with W155, W158, and Y383 (Fig. S10). In both upward-facing conformations, EOL’s two oxygen atoms are positioned near S151, facilitating hydrogen bond formation, a feature unique to EOL, as DEET lacks comparable oxygen atoms.

By contrast, DEET’s carbonyl oxygen in the upward-facing orientation points toward the top of the pocket, where it can form a hydrogen bond with surrounding water molecules. These water molecules, in turn, can interact with the backbone of E205 and A206, establishing a water-mediated bridge between DEET and the protein. This network effectively couples DEET to the binding pocket in a manner unavailable to EOL, which lacks both exposed oxygen atoms and the capacity for water-mediated hydrogen bonding. Such interactions may contribute to DEET’s binding efficiency, highlighting the role of solvent-mediated interactions in ligand recognition.

Additionally, EOL’s unique *∼*125*^◦^* orientation (State 3) corresponds to a “downward-facing” conformation, where the ligand primarily interacts with Y383 rather than W155 and W158 (Fig. S10). These structural observations align with the statistical analysis of ligand–pocket contact frequencies (Fig. 1c), demonstrating that EOL adopts multiple binding postures to engage distinct residues. Unlike DEET, which relies primarily on W155, W158 and water-mediated interactions, EOL forms more frequent van der Waals contacts with Y383 while reducing interactions with W155 and W158. Together, these findings indicate that EOL and DEET employ complementary binding strategies: EOL through direct residue contacts across multiple orientations, and DEET through solvent-mediated stabilization—potentially explaining differences in their binding efficiency and specificity.

Overall, the unbiased MD simulations of MhOR5 in complex with EOL and DEET revealed that both ligands were recognized by extensive van der Waals interactions with the pocket, with the most frequently contacted residues being largely similar. EOL has a unique hydrogen bond interaction with S151 and demonstrates two distinct orientations in the pocket—one facing upwards and the other downwards. In contrast, DEET consistently preferred one upward orientation.

### Ligand-Induced S4 Helix Kinking Underlies MhOR5 Binding Specificity

Building on binding pose analyses that demonstrated the superior stability of DEET-bound MhOR5 relative to its EOL-bound counterpart, we next conducted a comprehensive contact analysis to delineate how these divergent ligand-binding modes regulate the conformational dynamics of MhOR5. Our findings showed that, as mentioned above, unlike EOL, DEET forms an additional interaction outside the canonical binding pocket: a water-mediated hydrogen bond with residues E205 and A206 in the S4 helix (Fig. 2a). Specifically, the hydrophilic oxygen of DEET in its upward-facing pose engages water, which in turn hydrogenbonds to stabilize itself. This water-mediated hydrogen bonding interaction effectively couples DEET to S4 and is absent in the EOL-bound and apo systems (Fig. S9). Furthermore, we found that due to the upward-facing hydrophilic groups of DEET, there are significantly more water molecules in the binding pocket when DEET is bound compared to when EOL is bound and apo (Fig. 2a).

**Figure 2:**
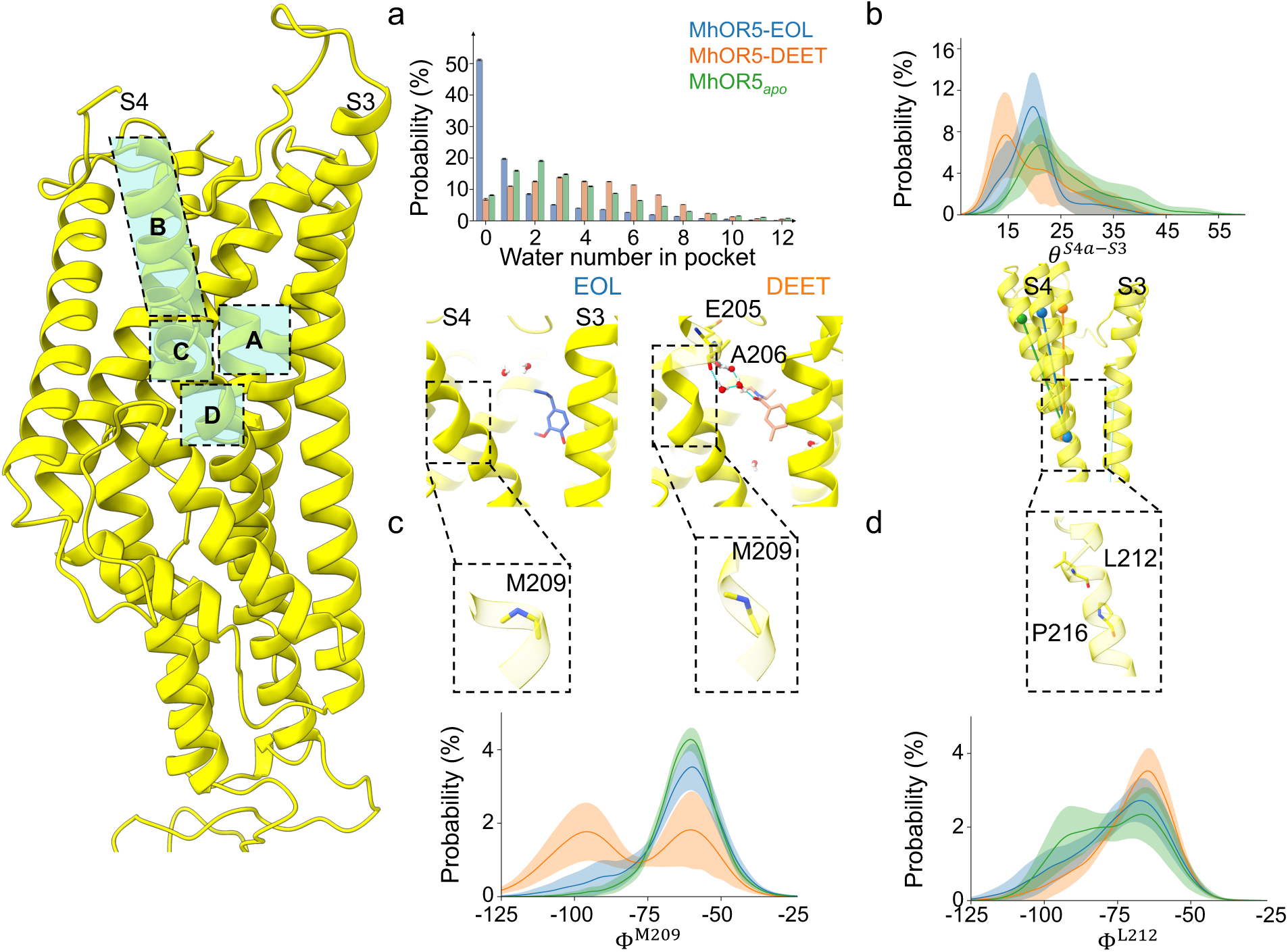
Odorant binding induces distinct S4 helix dynamics and binding pocket solvation in MhOR5 revealed by unbiased MD simulations. a) Histogram of water molecules within the ligand-binding pocket. DEET binding is associated with increased pocket hydration and formation of hydrogen bonds with residues E205 and A206, whereas such interactions are absent in the EOL-bound state. b) Probability distribution of the angle between S4a (upper segment of S4) and S3, revealing ligand-dependent dynamics of the transmembrane helices. c) Backbone dihedral angle (Φ and Ψ) distributions of residues M209. Upon DEET binding, an additional dominant, non-canonical conformation emerges, inducing a local kink in helix S4. d) Disruption of the hydrogen bond between P216 and L212 impairs the *α*-helical secondary structure of S4, leading to increased conformational dynamics, further modulating ligand specificity. All error bars in this figure are 95% CI calculated from each independent replica to show the variability between simulation runs explicitly.

To quantify changes in S4 conformation, we divided the helix into two segments—S4a (residues E199–S218) and S4b (residues T214-C236), and calculated the angle between each segment and the reference S3 helix, defined by the vector connecting the backbone centers of residues K134–F138 and G154–W158. S4b remained relatively stable across all systems, whereas S4a displayed pronounced ligand-dependent variability. The S4a angle was smallest in the DEET-bound state, intermediate in the EOL-bound state, and largest in the apo state (Fig. 2b). This hierarchy is consistent with the stabilizing interactions: DEET forms a persistent hydrogen-bond network that pulls S4 inward, EOL relies primarily on hydrophobic contacts without additional anchoring, and the apo state lacks any ligand-dependent stabilization.

To identify the structural origin of the S4a angle differences, we calculated residue-wise Φ*/*Ψ distributions for residues I203–I221. Four residues—A208, M209, L212, and I213—showed substantial deviations from canonical *α*-helical backbone dihedral angles. In A208 and M209, the DEET-bound ensemble exhibited two distinct Φ*/*Ψ populations: one matching the EOL and apo conformations, and another corresponding to a ligand-stabilized kink unique to DEET. This kink is consistent with the inward-bent S4a conformation observed only in the DEET-bound state (Fig. 2c).

Besides, L212 and I213 deviated from canonical angles, particularly in the EOL-bound and apo states, where their Φ / Ψ distributions were broader and slightly shifted (Figs. 2d and S11). This is attributable to a helix-breaker P216, which lacks a backbone amide hydrogen and disrupts the i→i+4 hydrogen bond in the helix (Figs. 2d and S11), inducing a local kink that affects residues L212–I213. In the DEET-bound state, however, a water-mediated hydrogen bond between DEET and S4 compensates for this disruption, stabilizing a more compact S4a configuration and narrowing the backbone dihedral distributions of L212–I213. Based on prior multiple sequence alignment (MSA) results,^29^ we focused on the allelic residue at position 216. Among 134 insect ORs, nearly half of the allelic residues were hydrophilic residues (Q or N) (Fig. S12). These hydrophilic residues located in the transmembrane hydrophobic region may also function to destabilize the helix, leading to universal S4 volatility.

Together, these findings establish a direct mechanistic link between ligand-specific binding interactions and the conformational plasticity of the MhOR5 S4 helix. DEET’s unique water-mediated hydrogen bonds with S4 enhance its binding stability relative to EOL, while inducing a localized S4a kink.

### Aromatic Ligands Share a Conserved Binding Mode in MhOR5

To assess whether the binding modes identified for EOL and DEET extend to related aromatic ligands, we examined 2-ethylphenol, 4-ethylphenol, and 4-methoxyphenylacetone, all of which are active against MhOR5 and share physicochemical features with EOL or DEET.^16^ All three ligands predominantly adopted the upward-facing orientation observed for EOL and DEET, supporting a conserved binding geometry (Fig. 3 and Figs. S13–S16). The two ethylphenols exhibited EOL-like binding patterns, including interactions with S151, whereas 4-methoxyphenylacetone showed a more restricted, DEET-like orientation with enhanced pocket hydration and weaker direct hydrogen interaction with S151 (Figs. S9 and S17).

**Figure 3:**
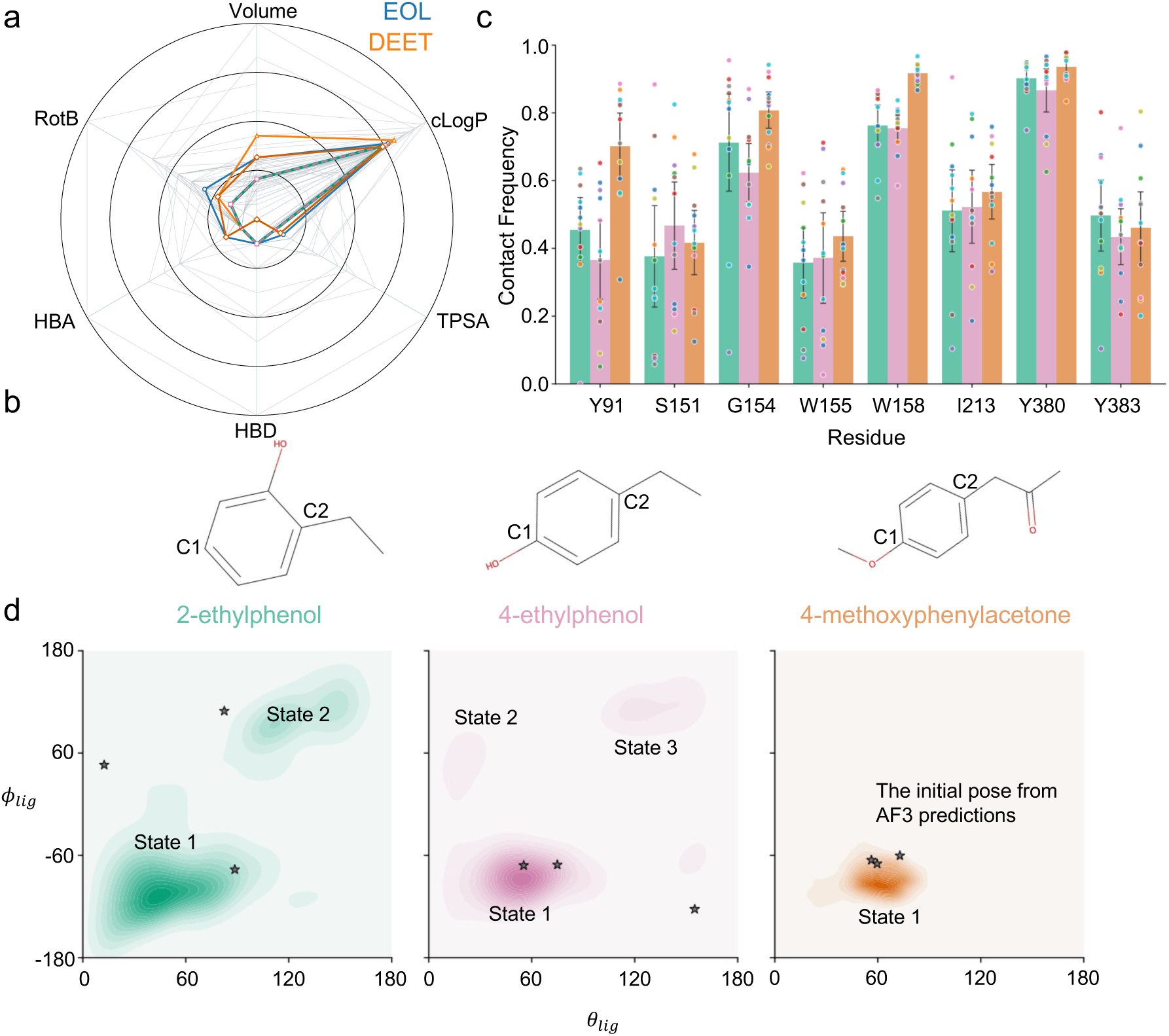
Physicochemical properties and binding modes of the expanded aromatic ligand panel. a) Radar plot comparing the molecular volume, calculated octanol– water partition coefficient (cLogP), topological polar surface area (TPSA), hydrogen-bond donor count (HBD), hydrogen-bond acceptor count (HBA), and rotatable-bond count (RotB) of compounds with reported electrophysiological responses against wild-type MhOR5. EOL and DEET are highlighted for reference. b) Two-dimensional structures of 2-ethylphenol, 4-ethylphenol, and 4-methoxyphenylacetone. C1 and C2 define the ligand long-axis vector used to calculate the orientation collective variables. c) Van der Waals contact frequencies between each ligand and eight binding-pocket residues. Bars show the mean across the 12 ligand–subunit trajectories, error bars denote trajectory-based confidence intervals, and colored points represent individual trajectories. d) Two-dimensional probability distributions of the ligand tilt angle, *θ_lig_*, and relative dihedral angle, *ϕ_lig_*, for the three ligands. Stars indicate the initial poses selected from AlphaFold 3 predictions, and the labeled states denote the major orientational ensembles sampled during the unbiased MD simulations.

These differences were accompanied by ligand-dependent S4 conformations. Two ethylphenols largely retained the EOL-like *θ^S^*^4*a−S*3^ distribution, whereas 4-methoxyphenylacetone shifted S4a toward the inward DEET-like conformation, consistent with its water-mediated hydrogen bonding (Figs. S9 and S17). Thus, the ligand-specific binding mode is compatible with distinct interactions and S4 responses.

Because quantitative kinetic and thermodynamic analyses require extensive sampling, subsequent calculations focused on EOL and DEET, for which ligand-bound MhOR5 cryoEM structures provide well-defined reference states.

### Spontaneous Ligand Binding to MhOR5: Kinetics and Pathways

After characterizing the atomic details of how aromatic ligands bind to MhOR5, we next investigated the dynamic processes and kinetics that underpin this binding. To facilitate the observation of binding events, we randomly placed 10 copies of either EOL or DEET in the solvent phase—at a distance from MhOR5’s binding pocket—resulting in a ligand concentration of *∼*9.6 mM. We then performed 10 independent simulations for each model, designated as MhOR5*_apo_*-10EOL or MhOR5*_apo_*-10DEET, each simulation run lasting up to 1000 ns.

Notably, we observed two spontaneous binding events for EOL and six for DEET during these simulations, from which we estimated the mean binding time *τ_on_* and the binding rate (*k_on_*) using the maximum likelihood estimation (MLE, see the Methods section for details).^30,31^ The results indicated that DEET’s binding rate (16.18 *±* 6.61 *×* 10^6^*M^−^*^1^*s^−^*^1^) was approximately three times faster than EOL’s (5.27 *±* 3.73 *×* 10^6^*M^−^*^1^*s^−^*^1^), as summarized in Table S3.

We analyzed the resulting binding poses in relation to the cryo-EM structure. For EOL, both binding events yielded a pose consistent with that in the experimental structure (*θ_lig_ ≈* 60*^◦^*; Figs. S18 and S19). For DEET, three of the six binding poses converged to the cryo-EM conformation.

Further analysis revealed that the binding process could be categorized into two primary pathways: (1) direct translocation from the solvent phase along the membrane normal (Pathway P1), and (2) lateral diffusion from the membrane interior (Pathway P2) (Fig. 4a, b). Pathway P2 was further subdivided into two subroutes: translocation between transmembrane helices S1 and S4 (P2.1) and between helices S3 and S4 (P2.2) (Fig. S20). Ligands traversing Pathway P1 maintained a high degree of hydration prior to entering the binding pocket. In contrast, ligands utilizing Pathway P2 underwent a marked reduction in hydration upon membrane penetration, with this low-hydration state persisting following entry into the binding pocket (Fig. S21).

**Figure 4:**
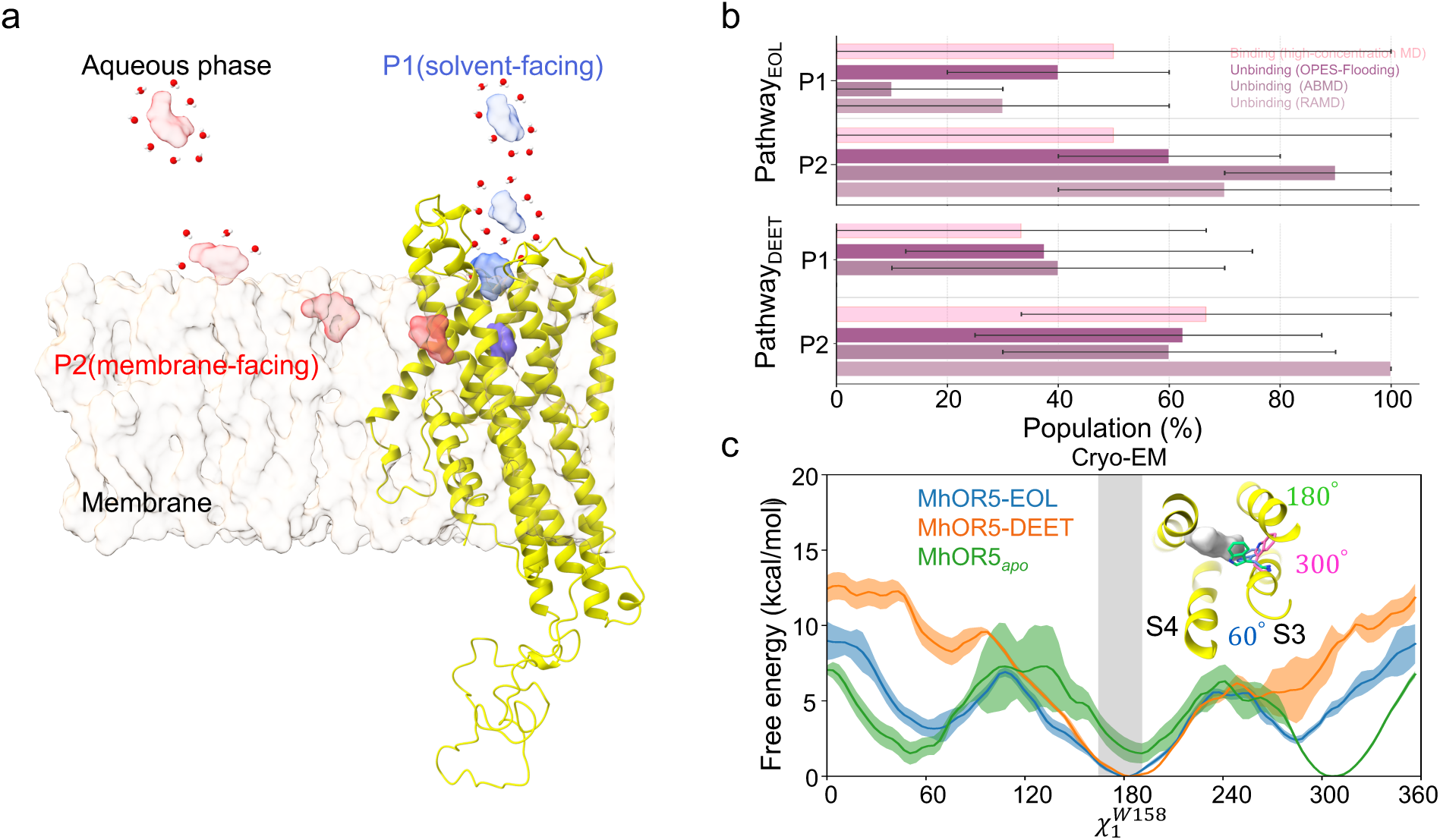
Two distinct pathways for ligand binding and dissociation. a) Two representative ligand-access pathways observed in simulations are depicted: P1 (blue), which faces the solvent and is exposed to the aqueous phase, and P2 (red), which faces the membrane and lies buried against the lipid bilayer. Red spheres indicate water molecules surrounding the hydrated ligand. b) Population distribution of EOL and DEET along the two pathways, derived from high-concentration spontaneous binding simulations and three dissociation approaches: RAMD, ABMD, and OPES-Flooding. Error bars represent the 95% CI estimated via 1,000 bootstrap resamples. c) Free energy landscapes of the W158 *χ*_1_ dihedral angle (*χ^W^* ^158^) in MhOR5–EOL, MhOR5–DEET, and MhOR5*_apo_* systems. The uncertainties 95% CI were obtained using block analysis with 10 blocks; solid lines indicate mean free energy values and shaded regions denote the associated uncertainties. The gray shaded area marks the range of *χ^W^* ^158^ angles observed in the corresponding cryo-EM structures. Inset top-view representations illustrate three representative *χ^W^* ^158^ conformations corresponding to open and closed states of the binding pocket. The ligand is shown in gray, and the W158 sidechain with different orientations is rendered as sticks in distinct colors.

### Enhanced Sampling Reveals Ligand Dissociation Pathways and the Gating Role of W158 in MhOR5

Our initial unbiased MD simulations of the ligand-bound MhOR5, limited to the microsecond timescale, did not capture dissociation events. To overcome this limitation, we employed two independent enhanced sampling techniques: random acceleration molecular dynamics (RAMD) and adiabatic bias MD (ABMD), for cross-validation. RAMD applies a randomly oriented force to the ligand to facilitate its unbinding, ^24^ while ABMD uses a ratchet-like biasing potential to accelerate the transition from the binding state to the unbinding state. ^25^ We performed ten simulations using each method for both ligand-bound states.

Consistent with spontaneous binding simulations, RAMD and ABMD simulations identified the same two primary dissociation pathways—P1 (toward the solvent) and P2 (toward the membrane) (Fig. 4a, b)—with a preference for P2-mediated exit. In all dissociation events captured by RAMD and ABMD, ligands were observed to first traverse the W158 residue prior to complete dissociation. This observation suggests that W158 may function as a gate regulating ligand release from the binding pocket.

### KTICA Identifies Key Dynamic Modes in Dissociation

To validate the potential gating function of W158 as a kinetic barrier, an adaptive sampling strategy based on macrostate counts was employed (see the Methods section for details). This strategy not only reduces bias toward spontaneous unbinding pathways but also improves sampling efficiency. In the W158-crossing events captured by the MD trajectories of adaptive sampling, W158 exhibited significant rotational motion. This motion was characterized by a *χ*_1_ dihedral angle (*χ^W^* ^158^) ranging from 50*^◦^* to 80*^◦^* (Fig. S22). Additionally, two dissociation events were observed in the 24 adaptive sampling MD trajectories; in both cases, the ligand itself had already passed beyond W158, and these events followed the P2 pathway.

To characterize the slow dynamic modes underlying these dissociation events, we employed the Koopman-reweighted time-lagged independent component analysis (KTICA) machine learning algorithm. Focusing on the binding pocket, we selected 120 candidate features for KTICA (see Methods for details). To quantify the dissociation process, we defined several ligand-residue distances and their negative exponent power, including *d_l−p_*, representing the distance between the ligand and the pocket center. And to quantify pocket dynamics, we considered the sine and cosine values of all dihedrals of the binding pocket residues. As a result, two leading components, KTIC 1 and KTIC 2, were identified.

Weight analysis of these KTICs revealed that the cosine function of *χ^W^* ^158^ and the sine function of *χ^W^* ^158^ were the top two most significant variables in KTIC1 (Fig. S23). And in KTIC2, the distance terms related to ligand dissociation account for the majority. Thus, KTIC1 primarily encompasses information regarding pocket dynamics (driven by W158 rotation), while KTIC2 reflects the ligand dissociation process.

To elucidate the features encoded in KTIC1 and comprehensively characterize the sidechain dynamics of W158, we employed the on-the-fly probability-enhanced sampling (OPES) algorithm to explore the free energy landscape of W158 sidechain dynamics in both apo and ligand-bound states, using *χ^W^* ^158^ and *χ^W^* ^158^ as CVs. The FES showed that in both ligand-bound systems, W158 primarily adopts a “gate-closed” state (consistent with the cryo-EM structure), which blocks ligand exit. A minor “gate-open” state was also observed, characterized by a shallower free energy basin where *χ^W^* ^158^ ranges from 240*^◦^* to 110*^◦^*, this open state facilitates ligand release (Fig. 4c, and Figs. S24-S25). Notably, the open state was less populated in MhOR5-DEET than in MhOR5-EOL, indicating that DEET encounters a higher energy barrier for pocket exit at this step. This is likely attributable to DEET’s more frequent direct interactions with W158 or water-mediated hydrogen bonding with the S4 helix. In contrast, the apo MhOR5 system exhibited three approximately equivalent free energy basins for *χ^W^* ^158^, indicating enhanced W158 dynamics and conformational heterogeneity in the absence of bound ligand.

These findings establish *χ^W^* ^158^ as a key CV linked to ligand dissociation. In contrast, analysis of *χ^W^* ^158^ revealed no substantial differences between the apo and ligand-bound states (Fig. S26), and ligand dissociation is independent of it (Figs. S27-S28), indicating it plays a less important role than *χ^W^* ^158^ in regulating the gating mechanism.

Taken together, these findings provide a mechanistic rationale for designating *χ^W^* ^158^ as the primary CV to further characterize W158-mediated ligand release in the subsequent section.

### Kinetics estimation of two release pathways

Leveraging the CVs defined through KTICA analysis, we employed the OPES-Flooding method with *d_l−p_* and *χ^W^* ^158^ as CVs to accelerate the unbinding kinetics of EOL and DEET and quantitatively calculate their dissociation rates.^26^

We conducted 20 OPES-Flooding simulations for each ligand-bound complex: simulations of MhOR5-EOL lasted up to 100 ns, while those of MhOR5-DEET were extended to 400 ns. This prolonged duration for DEET was justified by the W158 free energy landscape (FEL) derived from OPES, which indicated a potentially higher energy barrier for DEET release. All 20 MhOR5-EOL simulations yielded successful EOL dissociation; in contrast, only 8 of the 20 extended MhOR5-DEET simulations achieved DEET dissociation (Figs. S24-S25, S27-S28). A key observation was that *χ^W^* ^158^ undergoes rotational motion prior to ligand dissociation, which opens the binding pocket and enables ligand exit—indicating that ligands preferentially depart the deeply buried pocket only after W158 rotation (Figs. S24-S25). Consistent with the findings from ABMD and RAMD simulations, all dissociation events captured by OPES-Flooding occurred via both pathways P1 and P2.

These dissociation events allowed us to estimate the total mean dissociation/off time and off rate (*τ_off_* and *k_off_*) as well as pathway-specific values. We estimated *τ_off_* from a fit of the empirical cumulative distribution function (ECDF) to the theoretical cumulative distribution function (TCDF).^32^ The reliability of these estimates was evaluated a posteriori via the Kolmogorov–Smirnov (KS) test, where the computed p-value was used to verify the null hypothesis that transition time distributions follow a Poissonian model (Figs. S29 and S30). To account for unsuccessful dissociation events in our kinetic calculations and improve the comprehensiveness of our estimates, we additionally computed the total *τ_off_* and corresponding off-rates via MLE for MhOR5-DEET. Our results demonstrated that, relative to MLE-derived values, the KS test algorithm did not produce a significant overestimation of the total off-rate, yielding only an approximate twofold difference. This finding thus corroborates the reliability of kinetics estimates calculated by both methods, as summarized in Table 1.

**Table 1:**
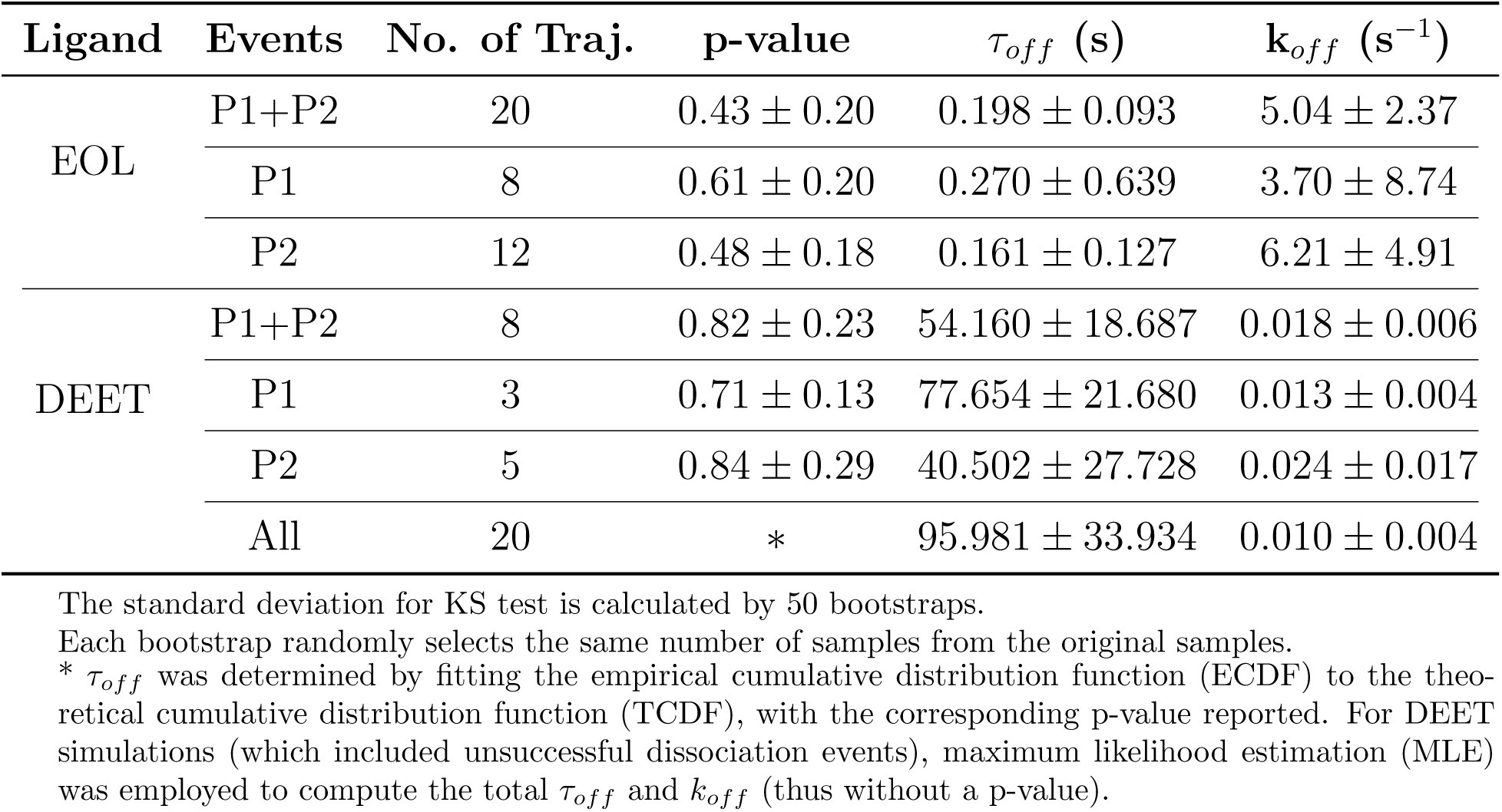
Dissociation times for EOL and DEET from the KS test and for DEET from Maximum Likelihood Estimation (MLE) of two pathways.

| Ligand | Events | No. of Traj. | p-value | $\tau_{off}$ (s) | $k_{off}$ ($s^{-1}$) |
| --- | --- | --- | --- | --- | --- |
| EOL | P1+P2 | 20 | $0.43 \pm 0.20$ | $0.198 \pm 0.093$ | $5.04 \pm 2.37$ |
| | P1 | 8 | $0.61 \pm 0.20$ | $0.270 \pm 0.639$ | $3.70 \pm 8.74$ |
| | P2 | 12 | $0.48 \pm 0.18$ | $0.161 \pm 0.127$ | $6.21 \pm 4.91$ |
| DEET | P1+P2 | 8 | $0.82 \pm 0.23$ | $54.160 \pm 18.687$ | $0.018 \pm 0.006$ |
| | P1 | 3 | $0.71 \pm 0.13$ | $77.654 \pm 21.680$ | $0.013 \pm 0.004$ |
| | P2 | 5 | $0.84 \pm 0.29$ | $40.502 \pm 27.728$ | $0.024 \pm 0.017$ |
| | All | 20 | * | $95.981 \pm 33.934$ | $0.010 \pm 0.004$ |

The standard deviation for KS test is calculated by 50 bootstraps.
Each bootstrap randomly selects the same number of samples from the original samples.
\* $\tau_{off}$ was determined by fitting the empirical cumulative distribution function (ECDF) to the theoretical cumulative distribution function (TCDF), with the corresponding p-value reported. For DEET simulations (which included unsuccessful dissociation events), maximum likelihood estimation (MLE) was employed to compute the total $\tau_{off}$ and $k_{off}$ (thus without a p-value).

Our findings revealed that EOL exhibits a significantly faster off-rate than DEET, with a difference of two to three orders of magnitude. Additionally, the off-time for EOL via the P2 (membrane-facing) pathway was *∼*160 ms, roughly half that of the P1 (solvent-facing) pathway; for DEET, the off-time via P2 was *∼*40 s, again approximately half that of P1 (Table 1). These results indicate that unbinding via the membrane pathway is slightly more favorable for both ligands.

In conclusion, the combination of enhanced sampling techniques and adaptive sampling provided a comprehensive understanding of the dissociation process. The whole ligand unbinding process can be divided into two steps. In the initial step, W158 undergoes rotational movement, allowing the ligand to exit the binding pocket and enter the vestibule, which is a significant difference from the binding process. In the second step, the ligand fully dissociates from the protein through two main pathways as same as the binding process.

### Calculations of odorant binding affinities

To accurately assess EOL/DEET binding affinities to MhOR5 and validate data consistency, we first calculated kinetic-based binding free energy 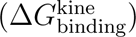 using prior kinetic data, then determined absolute binding free energy (ABFE) via FEP, followed by cross-validation of the two datasets.

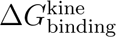 was computed using the association rate constant (*k*_on_) and dissociation rate constant (*k*_off_) derived from OPES-Flooding kinetic analyses, employing the equation: 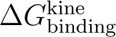 = *k_B_T* ln(*k_off_ /*[*k_on_C*_0_]). Here, *k_B_* denotes the Boltzmann constant, *T* is the absolute temperature, and *C*_0_ = 1 M represents the standard reference concentration. This approach directly correlates binding affinity with the kinetic processes of ligand binding and unbinding. ABFE was quantified via FEP with a thermodynamic cycle (Fig. S31a). Convergence was confirmed for both complex and ligand-only systems (Fig. S31b), ensuring reliable FEP results which showed DEET has a stronger affinity than EOL (Table S5). Cross-validation 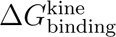 and 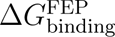 (Table 2) showed good agreement: MhOR5-EOL (–8.54 *±* 0.52 vs. –8.39 *±* 0.22 kcal/mol) and MhOR5-DEET (–13.06 *±* 0.34 vs. –11.25 *±* 0.16 kcal/mol, with a modest *∼*2 kcal/mol discrepancy). This discrepancy may arise from OPES-Flooding overestimation of slow kinetics or MLE sensitivity to outliers. Both datasets consistently confirmed DEET’s higher binding affinity.

**Table 2:** **Comparison of binding free energy, *k_on_*, and *k_off_***

| Ligand | $k_{\text{on}}(10^6 M^{-1} s^{-1})$ | $k_{\text{off}}(s^{-1})$ | $\Delta G_{\text{binding}}^{\text{FEP}}(\text{kcal/mol})$ | $\Delta G_{\text{binding}}^{\text{kin}}(\text{kcal/mol})$ |
| --- | --- | --- | --- | --- |
| EOL | $5.27 \pm 3.73$ | $5.04 \pm 2.37$ | $-8.39 \pm 0.22$ | $-8.54 \pm 0.52$ |
| DEET | $16.18 \pm 6.61$ | $0.010 \pm 0.004$ | $-11.25 \pm 0.16$ | $-13.06 \pm 0.34$ |

Note: $\Delta G_{\text{binding}}^{\text{FEP}}$ indicates that it is obtained from FEP simulation, and $\Delta G_{\text{binding}}^{\text{kin}}$ indicates that it is calculated from $k_{\text{on}}$ and $k_{\text{off}}$ .

Consistency between FEP and kinetic-based results validates the robustness of both thermodynamics and kinetics calculations, demonstrating DEET’s superior affinity stems from faster *k*_on_ and substantially slower *k*_off_ than EOL, aligning with earlier kinetic observations.

## Discussions and Conclusions

Recent advances in structural biology have enhanced our understanding of how insect ORs detect volatile molecules; however, critical questions persist regarding the mechanisms underlying ligand selectivity and entry/exit.^16–18,20,33^ The present study addresses these uncertainties by investigating the binding and unbinding dynamics of MhOR5, the earliest identified insect OR, using an integrated computational workflow.

Analysis of key residues elucidated the functional mechanisms of MhOR5: W158 was identified to play dual roles in receptor selectivity and gating ligand release, while W155 and Y383 were also found to contribute to receptor selectivity. Experimental mutagenesis studies—where Y383A and W158A mutations significantly attenuated receptor responses^16^—directly validate these computational findings, reinforcing the functional significance of W158 as a critical gating mechanism controlling ligand release (Fig. 5). This conclusion is further supported by recent computational work highlighting analogous gating roles of conserved residues in other ORs.^34,35^

**Figure 5:**
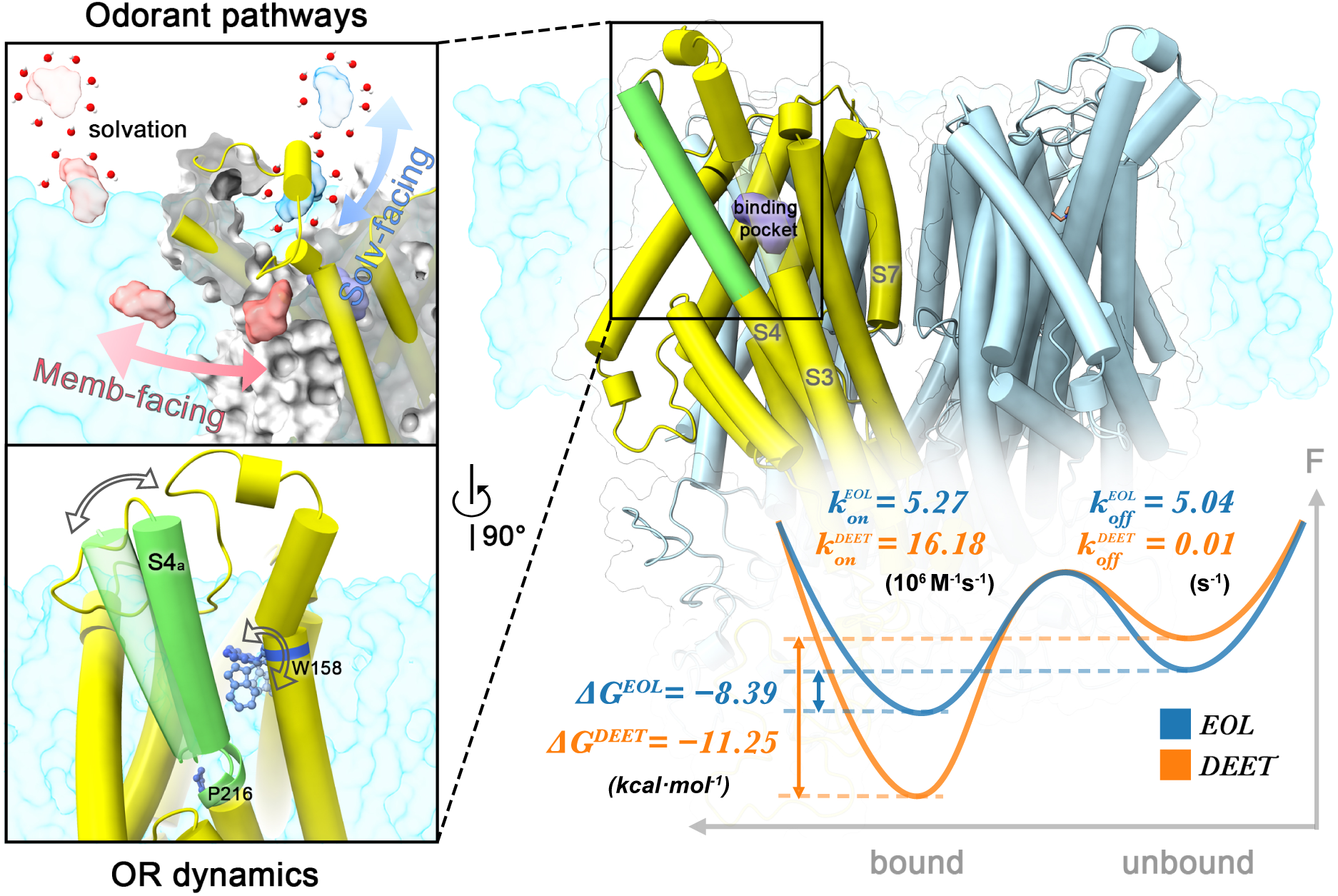
Integrated mechanism of ligand binding, gating, and dissociation in MhOR5. The upper-left panel illustrates two ligand entry and exit pathways: a solvent-facing pathway (P1) and a membrane-facing pathway (P2). The lower-left panel highlights ligand-induced structural rearrangements in helix S4, showing that P216 disrupts the local *α*-helical structure and increases S4 flexibility, while W158 is positioned at the binding pocket and is closely associated with ligand dissociation. The central and right panels depict the binding free energies and on/off rate for EOL and DEET, illustrating ligand-dependent differences in binding affinity and association/dissociation kinetics.

Furthermore, we identified P216 as a key contributor to the flexibility of the S4 helix. Sequence alignment analysis revealed that nearly half of 134 insect ORs harbor hydrophilic residues in the transmembrane domain at this position, suggesting that destabilization of helical secondary structure might be a conserved feature facilitating S4 helix conformational fluctuations. Different ligands induce distinct conformations of the S4 helix due to their varying binding modes: DEET’s water-mediated hydrogen bond network stabilizes a kinked, inward-bent S4a segment, enhancing binding stability, while EOL’s lack of such solvent-mediated anchoring results in a more flexible, outward-bent S4a. The observation that three additional aromatic ligands preserve this binding mode supports its broader relevance and provides a useful framework for rational selectivity-ligand design.

These findings were further supported by experimental evidence:^16^ the M209V mutation reduced responsiveness to EOL while enhancing sensitivity to DEET, likely due to weakened hydrophobic interactions with EOL and facilitated formation of water-mediated hydrogen bonds with DEET. In contrast, the I213M mutation reinforced hydrophobic interactions, thereby suppressing water-mediated hydrogen bonding and leading to the opposite functional outcome. This ligand-induced structural plasticity of the S4 helix may serve as a general mechanism for tuning OR binding affinity and specificity across insect species. This framework also helps rationalize the mutant responses of three related ligands: M209V selectively enhances 4-methoxyphenylacetone, consistent with its DEET-like polarity and flexibility, whereas I213M distinguishes eugenol from the two smaller ethylphenols, suggesting ligand-specific packing.

By integrating enhanced and adaptive sampling strategies, we achieved a comprehensive understanding of the ligand binding and unbinding process. The ligand dissociation proceeds in two steps: the first step involves the rotation of W158 enabling ligands to exit the binding pocket and enter the vestibule. In the second step, ligands undergo complete dissociation via two pathways shared with the binding process: one through the aqueous phase above the receptor and the other at the membrane interface. This membrane-mediated pathway preference, also reported for other systems,^36–38^ was observed for both lipid-soluble ligands, DEET and EOL. We note, however, that the number of binding events sampled in our simulations remains limited; therefore, a statistically robust quantification of pathway preference during the association process will require further sampling.

These findings support a proposed “two-step” dissociation model, which may be broadly conserved across insect ORs. A key dynamic feature underpinning this model is the distinct behavior of W158. In the apo state, W158 is intrinsically flexible and frequently samples an open conformation, permitting rapid ligand binding with no significant kinetic barrier. In contrast, during dissociation, the rotation of W158 from its closed (ligand-bound) to open state presents the major rate-limiting barrier for ligand exit.

To quantify ligand affinity, we calculated absolute binding free energies using FEP. The FEP-derived binding free energies, together with those estimated from the kinetic parameters of association rates (*k_on_*) and dissociation rates (*k_off_*), collectively confirm that DEET binds more tightly to MhOR5 than EOL. This finding is consistent with DEET’s water mediated hydrogen bonding interactions with the S4 helix and its more persistent van der Waals contacts with key residues (W155, W158). Notably, a minor discrepancy was observed in the DEET-related binding free energy values between the two approaches. This discrepancy is likely attributable to challenges in accurately estimating the slow ligand dissociation kinetics from biased OPES-Flooding simulations, as aggressive biasing may disrupt transition state regions and complicate the kinetic reweighting process. Furthermore, since not all trajectories underwent dissociation, we used MLE for estimation, which, compared to the KS test, makes the results sensitive to outliers. Overall, the close agreement between FEP and kinetic-based affinity estimates underscores the reliability of both methods.

Interestingly, DEET’s stronger binding affinity, as evidenced by the simulations, did not correlate with greater ligand-stimulated current intensity in experimental measurements.^16^ This discrepancy likely reflects the complex interplay between ligand binding and signal transduction. Binding affinity alone might be insufficient to predict functional responses, as downstream processes may be differentially modulated by ligand structure. It cannot be ruled out that part of this difference may also arise from force field inaccuracies.

Collectively, this work provides atomistic insights into the ligand recognition, binding, and release mechanisms of MhOR5, an ancestral insect OR. Our findings highlight that insect odorant responses are shaped not only by binding affinity but also by the kinetic and dynamic properties of ligand-receptor interactions—including binding/unbinding rates, pathway preferences, and ligand-induced conformational changes (Fig. 5). The identification of W158 as a gating residue, the characterization of dual binding/dissociation pathways, and the demonstration of S4 helix plasticity offer a framework for understanding OR function across insect lineages. Furthermore, the integrated computational approach employed in this study—combining unbiased MD simulations, enhanced sampling, machine learning, and FEP—serves as a robust template for investigating other chemoreceptors, including GRs and IRs, which share structural and functional similarities with ORs.

Future research should prioritize validating the proposed gating mechanisms and ligand transport pathways through complementary structural and functional experimental approaches. Additionally, extending this computational and functional analysis to OR-Orco heterocomplexes is essential, as the Orco co-receptor is a critical component of the functional OR complex in most insect species. Beyond elucidating receptor biology, integrating atomistic simulations with protein engineering may enable the development of next-generation OR-based biosensors and bioelectronic devices for sensitive detection of volatile compounds.^39^ This synergistic approach would enable the rational design and rapid evaluation of compounds targeting key residues or ligand pathways, thereby advancing the development of sustainable, species-specific pest management strategies while also broadening the technological applications of insect ORs.

## Methods

### Modeling of the apo and ligand-bound states of MhOR5

The cryo-EM structures of MhOR5 in its apo state (MhOR5*_apo_*), and its complexes with EOL (MhOR5-EOL) and DEET (MhOR5-DEET) (PDB IDs: 7LIC, 7LID, and 7LIG),^16^ were used as the templates to build the atomistic models. Missing loop regions were modeled using the SWISS-MODEL web server.^40^

These models were then embedded in a flat lipid bilayer made of 1-palmitoyl-2-oleoyl-sn-glycero-3-phosphocholine (POPC) and solvated in a cubic water box containing 0.15 M NaCl. The water box dimensions were 12.0 nm × 12.0 nm × 15.8 nm, resulting in approximately 210,000 atoms per system. The PPM web server (version 2.0) was used to align the protein within the lipid bilayer.^41^ System construction was carried out using the CHARMM-GUI web server,^42^ with both the N-terminal and C-terminal patched by neutral patches (NNEU and CNEU) to maintain electrical neutrality. The protonation states of all titratable residues were determined at pH 7.0 using PROPKA 3.0 via the CHARMM-GUI platform.^43^ Unless otherwise specified, all residues were maintained in their standard protonation states, and histidine residues were modeled in the tautomeric form with protonation at the *N_δ_* position.

The CHARMM36m force field was applied to the proteins,^44^ and the TIP3P model was used for water.^45^ The CHARMM-WYF parameters were utilized for cation-*π* interactions, and the ligands were parameterized using the CGenFF force field. ^46^

In the MD simulations, the temperature was kept constant at 310 K using a V-rescale thermostat with a 1 ps coupling constant, and the pressure was kept at 1.0 bar using the Parrinello–Rahman barostat with a 5ps coupling constant. A cut-off of 1.2 nm was applied for the van der Waals interactions using a switch function starting at 1.0 nm. The cut-off for the short-range electrostatic interactions was also set at 1.2 nm, and the long-range electrostatic interactions were calculated using the particle mesh Ewald decomposition algorithm with a mesh spacing of 0.12 nm. A reciprocal grid of 96 × 96 × 129 cells was used with fourth-order B-spline interpolation. All simulations were performed using a GPU-accelerated version of GROMACS 2023.^47^

### Unbiased MD simulations and data analysis

Following an initial energy minimization using the steepest descent algorithm, a six-step equilibration process was performed, gradually releasing positional constraints. Production MD simulations were then conducted under semi-isothermal-isobaric (NPT) conditions. The hydrogen mass repartitioning technique (HMR) ^48^ was applied, yet a 2 fs time step was implemented for stability. To assess the potential impact of HMR on system dynamics and kinetics, we conducted a baseline study using an alanine dipeptide (Ala2) as the model system (see SI for details). Each simulation was run for 1000 ns. The MSM analysis confirms that using HMR with a 2 fs time step does not alter the thermodynamics and kinetics of the system (Fig. S32).

Interactions between ligands and the protein were analyzed using the open-source Python package GetContacts.^49^ Root Mean Square Deviation (RMSD) was performed using Gromacs tools.

To investigate the orientation dynamics of EOL and DEET, we projected the trajectories onto two CVs: the tilt angle *θ_lig_* and the dihedral angle *ϕ_lig_*, whose definitions are detailed below.

- *è_lig_* (Tilt Angle): Defined as the angle between each ligand’s long axis *r_C_*^1^−*_C_*^2^ and the membrane normal *r_Dum_*_1_*_−Dum_*_2_, reflecting the ligand’s tilt relative to the membrane normal. For EOL, *r_C_*^1^−*_C_*^2^ extends from the benzene carbon *C*^1^ (attached to the hydroxyl group) to the benzene carbon *C*^2^ (linked to the allyl group) (Fig. 1d). For DEET, it runs from the para carbon *C*^1^ of the benzene ring to the carbonyl carbon *C*^2^ of the amide group. And *r_Dum_*_1_*_−Dum_*_2_ is ion channel vector from the center of ion channel to the point where the center of the ion channel is shifted along the membrane normal.
- *ϕ_lig_* (Dihedral Angle): A relative dihedral angle, defined as the angle between two planes: (1) plane *n*_(*r*_*_C_*_1_*_−C_*_2_ *_,rDum_*_1_*_−Dum_*_2)_, formed by the ligand vector *r_C_*^1^−*_C_*^2^ and ion channel vector *r_Dum_*_1_*_−Dum_*_2_ ; (2) plane *n_C_*^1^*_−Dum_*_1_*_−Dum_*_2_, formed by the ion channel vector *r_Dum−Dum_* and the ligand atom *C*^1^. This dihedral provides a monomer-independent orientation measure: *ϕ_lig_* = 0 indicates the ligand points directly toward the central ion channel, while values approaching *±*180*^◦^* indicate it points away from the pore.

Ligand orientation, S4 helix dynamics and related dihedral angles were performed using the PLUMED driver function (version 2.9).^50^ The number of water molecules in the pocket was performed using MDAnalysis package.^51^ Probability distributions were computed using gaussian kernel density estimation (KDE) algorithm in the SciPy package.^52^ The sigma value is selected to be one in a thousand of the entire CV space, i.e. 0.00314. Error bars are calculated based on independent replicas.

### Simulations of expanded aromatic ligands

To extend the chemical space beyond EOL and DEET while retaining comparable physicochemical properties, we first collected ligands with reported electrophysiological responses against MhOR5 and calculated their molecular volume, cLogP, topological polar surface area (TPSA), hydrogen-bond donor (HBD) and acceptor (HBA) counts, and rotatable-bond (RotB) counts using RDKit.^16^ Aromatic compounds with physicochemical properties close to either EOL or DEET were retained, leading to the selection of 2-ethylphenol, 4-ethylphenol, and 4-methoxyphenylacetone (Fig. 3a). For each ligand, 500 AlphaFold 3 models were generated from 100 random seeds, with five samples per seed. Models in which the ligand heavy-atom centroid was located within 3 Å of an EOL-binding site were retained. The retained ligand poses were then clustered by average-linkage hierarchical clustering based on heavy-atom RMSD, and the medoid of each of three clusters was used as an independent starting structure.

Each medoid pose was placed in the four equivalent binding pockets of the MhOR5 tetramer. System construction and unbiased MD simulations followed the same protocol used for EOL and DEET. After energy minimization and the six-step restrained equilibration, three independent 1 *µ*s production simulations were performed for each ligand at 310 K. Because each MhOR5 tetramer contains four equivalent subunits, the three simulations provided 12 ligand–subunit trajectories per compound. To investigate the orientation dynamics of the three ligands, we projected the trajectories onto the tilt angle *θ_lig_* and relative dihedral angle *ϕ_lig_*, following the same definitions used for EOL and DEET. The ligand long axis was defined by the vector connecting C1 and C2 indicated in Fig. 3b. Ligand orientation, S4 helix dynamics, and the related backbone dihedral angles were calculated using the PLUMED driver function.^50^ Interactions between ligands and the protein were analyzed using GetContacts.^49^ Pocket water numbers were calculated with MDAnalysis,^51^ and probability distributions and trajectory-based confidence intervals were obtained as described for EOL and DEET.

### Ligand binding pathways and on-rate estimations from unbiased MD simulations

To investigate ligand binding pathways and on-rates, we adopted an unbiased MD approach, aiming to observe spontaneous ligand binding. For both EOL and DEET, the ’gmx insert-molecules’ command was used to randomly place 10 identical ligands in the simulation box of apo MhOR5 (PDB ID: 7LIC), followed by 1000 ns of unbiased MD. To accommodate the added ligands and prevent self-aggregation due to high local concentration, the simulation box was enlarged to dimensions of 15.0 nm × 15.0 nm × 13.4 nm. Ten parallel simulations were conducted for each model, resulting in a total of 20 simulations.

The effective ligand concentration in the simulation was approximately 9.6 mM, calculated as: [*L*] = 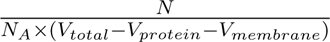 where N=10 is the number of ligand molecules, *N_A_* = 6.02 *×* 10^23^*mol^−^*^1^ is the Avogadro constant, *V_total_ ≈* 3.0 *×* 10^3^nm^3^ is the total box volume, *V_protein_ ≈* 2.5 *×* 10^2^ nm^3^ is the protein volume (calculated using the volume tools in ChimeraX^53^), and *V_membrane_ ≈* 1.0 *×* 10^3^nm^3^ is the volume occupied by the membrane slab.

Given that MhOR5 is a homotetramer, with a maximum of two binding events observed in a single simulation. The ligand association rate constant (*k_on_*) and its corresponding uncertainty were computed using maximum likelihood estimation (MLE),^30,31^ with the specific calculation equations provided below (a detailed derivation of the MLE framework is available in the Supplementary Information):

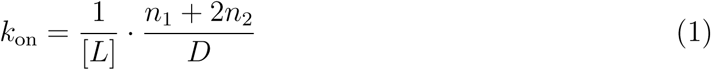

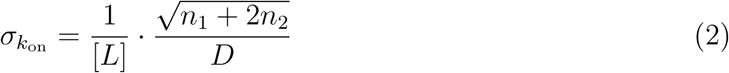

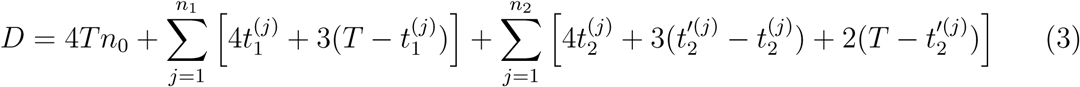

In the above equations:

- *T* : Total duration of each simulation (uniform across all replicates),
- *n*_0_: Number of simulations with no observed binding events,
- *n*_1_: Number of simulations with one observed binding event, where 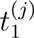 denotes the binding time in the *j*-th such simulation,
- *n*_2_: Number of simulations with two observed binding events, where 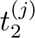 is the time of the first binding event and 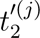 is the time of the second binding event in the *j*-th such simulation,
- [*L*]: Concentration of the ligand used in the experiments.

Further calculations (as outlined in SI) include: The ligand on-time (*τ*_on_), computed as:

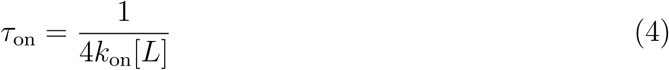

The dissociation constant (*K_d_*), derived from the relationship between association and dissociation rate constants:

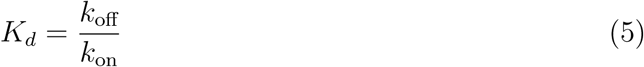

where *k*_off_ is the ligand dissociation rate constant, The binding free energy (Δ*G*_binding_), calculated using the Boltzmann distribution:

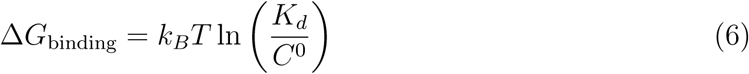

where *k_B_* is the Boltzmann constant, *T* is the absolute temperature, and *C*^0^ is the standard reference concentration (fixed at 1 M).

The uncertainties associated with *τ*_on_, *K_d_*, and Δ*G*_binding_ were determined via error propagation analysis, as specified in SI.

### Random acceleration MD (RAMD) and Adiabatic-bias MD (ABMD)

To accelerate the ligand unbinding process, we utilized RAMD^24^ and ABMD,^25^ which were conducted using the same simulation parameters as unbiased MD, but included additional bias forces or potentials.

In RAMD, a randomly oriented force facilitates the ligand unbinding. This force is reassigned randomly if the substrate’s movement falls below a set threshold distance within a specified time interval. The reference pocket for applying the bias was defined using the center of mass of Y91, S151, G154, W155, W158, I213, Y380, and Y383. The force constant was set at 160 *kJ/*(*mol · nm*^2^) with a maximum ligand-reference distance of 10 nm. The simulations were performed using GROMACS-RAMD.^54^

ABMD employs a ratchet-like biasing harmonic potential, which remains zero as the system moves towards the unbound state and increases when moving backward to the bound state. The ABMD simulations were carried out using the PLUMED 2.9,^50^ with the target point set at 5.0 nm and the strength of the bias (KAPPA) at 24 *kJ/*(*mol · nm*^2^).

### Adaptive sampling strategy guided by Koopman-reweighted TICA

TICA (time-lagged independent component analysis) is an unsupervised machine learning method for identifying slow order parameters.^55,56^ TICA enhanced by Koopman reweighting (KTICA) uses importance sampling to derive weight functions for each time-lagged pair, avoiding symmetry enforcement biases caused by statistical noise or insufficient sampling.^57^ Adaptive sampling accelerates conformational space exploration by selecting seed states from short unbiased trajectories^58–60^ and proceeding in round-by-round incremental steps.

Candidate features input into KTICA and Markov State Models (MSM) included ligand dissociation behavior and pocket dynamics. Ligand dissociation was quantified by the distance between the ligand and the aforementioned pocket residues and their negative exponent power (see Table S4 for details). The dynamics of the pocket were quantified by the sine and cosine values of all dihedral angles of these residues.

The lag time for KTICA and MSM was set to half the length of the short trajectory (i.e., 25 ns). The Deeptime package was used to perform KTICA analysis and construct the MSM.^61^ After feature transformation and dimensionality reduction via KTICA, microstates were obtained via clustering with the minibatch k-means algorithm. ^62^ The PCCA+ (Perron Cluster Cluster Analysis) algorithm, which further groups microstates based on their pairwise kinetic distances, was applied to construct macrostates.^63^

First, a single trajectory derived from prior ABMD simulations—where the complete ligand dissociation process spanned 52 ns—was selected and partitioned into 52 structural snapshots. Given that the ligand only underwent local fluctuations within the binding pocket during the initial 40 ns of the trajectory, the final 12 ns trajectory, which shows the dissociation process, was extracted. From this trajectory, structural snapshots were sampled at 1 ns intervals to serve as initial conformations for subsequent 50 ns simulations per snapshot. This setup yielded a cumulative trajectory length of 600 ns for the tetramer for the first round of adaptive sampling. Since the four subunits are equivalent, we effectively obtained a 2.4 *µs* trajectory for a single monomer. Based on macrostate counts,^64,65^ 12 new starting structures were selected from these 2.4 *µs* trajectories, each of which was subjected to an additional 50 ns simulation in the next sampling round. Following the generation of a combined dataset totaling 4.8 *µs* of trajectories, KTICA was implemented to identify linear combinations of candidate descriptors, designated as KTICA components.

Although KTICA components can obscure direct physical interpretation, the coefficient *b_i_* associated with each KTICA component serves as a measure of the relative importance of each input feature to that component. To quantify the contribution of the input features to the KTICA components, we performed the weight analysis by analyzing the corresponding expansion coefficients *b_i_*.

### OPES to explore the free energy landscape of W158 dynamics

The on-the-fly probability-enhanced sampling (OPES) algorithm was employed to explore the free energy landscape of the MhOR5-EOL, MhOR5-DEET, and MhOR5*_apo_* along the *χ*_1_ and *χ*_2_ of W158.^66^ The BARRIER parameter was set to 40 *kJ/mol*, and the frequency of adding bias was set to 1 ps. To focus on exploring the conformational space in the bound state, an UPPER WALLS potential was applied to *d_l−p_*at 1.2 nm, and the bias constant (KAPPA) was set to 150 *kJ/*(*mol· nm*^2^) to prevent ligand dissociation. The simulation time for MhOR5-EOL, MhOR5-DEET and MhOR5*_apo_* was set to 1000 ns. FESs were computed using the reweighting script available on the PLUMED tutorial website.^67^ 10 blocks are employed to calculate the error bar. The sigma value is selected to be one percent of the entire CV space, i.e. 0.0628.

### OPES-flooding to calculate the ligand off rates

In OPES-Flooding,^26^ a 2D bias was introduced along the *d_l−p_* (the distance between the ligand center and the pocket center) and the *χ^W^* ^158^. The OPES METAD EXPOLRE algorithm provided in PLUMED 2.9 was employed with the BARRIER parameter set to 20 *kJ/mol* and the bias frequency set to 1 ps. Bias was only added to the area where *d_l−p_* was less than 1.6 nm, to avoid interfering with the transition state. At this distance, the ligand has just crossed W158 but is far from dissociating. Once the distance exceeds 2.4 nm, the ligand has completely dissociated so that the simulation will be terminated. MhOR5-EOL and MhOR5-DEET each underwent 20 simulations, with durations of up to 100 ns or 400 ns, respectively.

Subsequently, the mean off-rate time (*τ_off_*) was estimated by fitting the empirical cumulative distribution function (ECDF) to the theoretical cumulative distribution function (TCDF).^32^ The reliability of these *τ_off_*estimates was validated post-hoc using the Kolmogorov–Smirnov (KS) test. Specifically, the computed p-value from the KS test was used to verify the null hypothesis that the transition time distributions conform to a Poissonian model.^26,68,69^ Error bars for *τ_off_* were determined via 50 bootstrap resampling iterations, where each iteration involved random sampling of an equal volume of data from the original dataset.

For DEET, successful dissociation events were not observed in all simulations; thus, MLE was again used to estimate total (without distinguishing between the different pathways) off-time.^30,31^ Since dissociation events only involve monomers, the equation is:

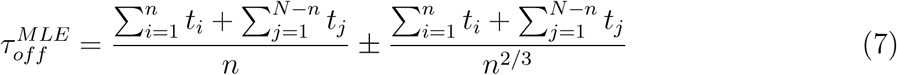

Here, *t_i_* denotes the dissociation time for each of the *n* successful dissociation events, while *t_j_* represents the duration of the remaining *N − n* trajectories in which dissociation had not occurred by the end of the simulation period.

The dissociation rate constant (*k_off_*) for all pathways was derived as the reciprocal of its corresponding *τ_off_* value. Uncertainties associated with *k_off_*were computed using the standard error propagation formula.

### FEP-based method to calculate the absolute binding free energy

PMX was used to create a hybrid topology that contains both vanish and emergence states for the alchemical transformations and imposing 6 restraints (1 distance, 2 angles, and 3 torsions).^70^ The strength of restraints are 3000 *kcal/mol/nm*^2^ for distance and 30 *kcal/mol/rad*^2^ for angles and torsions. The absolute binding free energy for MhOR5-ligand (Δ*G*_binding_) was determined using the thermodynamic cycle depicted in (Fig. S31a) in which ΔΔ*G*_binding_ = Δ*G*_3_ + Δ*G*_4_ *−* Δ*G*_1_.

For ligand annihilation in a ligand-only system, 21 lambda windows were used (10 ns per window) to ensure measurement accuracy. For ligand annihilation and restraint imposition in the complex system, we referenced the Lucid double-decoupling method (LDDM), gradually adding restraints and annihilating the force field across 18 windows to improve window overlap.^71^ Each window was simulated for 40–120 ns to account for varying window overlap difficulty.

Data post-processing involved removing correlated data, followed by application of the MBAR (Multistate Bennett Acceptance Ratio) algorithm via the Alchemlyb Python package.^72,73^

### Data visualization and representation

Trajectory and structure visualization was conducted using PyMOL. Plots were generated using Matplotlib 3.7 and Python 3.10.

## Data availability

The data that support this study are available from the corresponding authors upon request. The input files and scripts relating to the MD simulations can be found at GitHub [https://github.com/TiefengSong/MhOR5]. The raw MD trajectories are available at Zenodo 10.5281/zenodo.20119733].

## Supporting Information

The Supporting Information is available free of charge at [Pending upon publication].

Comprehensive methodological derivations, control simulations, and extended trajectory analyses that validate the computational framework, present additional structural and dynamic data, and detail the convergence of free energy calculations central to the study.

## Author Contributions

Y.W. conceived and supervised the project; T.S. performed all simulations and analyzed all simulations together with M.L. under the supervision of Y.W.; H.F. and H.S. participated in discussions during the study and contributed to the development of the methodology; T.S. wrote the first draft of the manuscript with input from Y.W. All authors contributed to the writing of the manuscript.

## Acknowledgement

We thank Mingyuan Zhang for assistance with writing and discussions. Y.W. acknowledges the financial support of the Zhejiang Provincial National Science Foundation of China (No. LZ24C050003), the National Natural Science Foundation of China (No. 32371300), and computational support of the Information Technology Center and State Key Lab of CAD&CG at Zhejiang University.

