## Supplementary material for "Ligand Binding Kinetics, Thermodynamics, and Gating of Insect Odorant Receptor": SI

### Supporting Information for: Ligand Binding Kinetics, Thermodynamics, and Gating of Insect Odorant Receptor

#### S1 Derivation of Maximum Likelihood Estimation (MLE) for Ligand Binding Kinetics

This section provides a detailed, step-by-step derivation of the Maximum Likelihood Estimation (MLE) for the ligand association rate constant $k_{\rm on}$ and its standard deviation $\sigma_{k_{\rm on}}$.

##### S1.1 Fundamental Assumptions

###### S1.1.1 Poisson Process and Exponential Distribution of Binding Events

Ligand binding to the tetrameric MhOR5 receptor follows a Poisson process. For such a process, the probability density function (PDF) of the time $t$ at which a binding event occurs obeys an exponential distribution:

$$f(t) = \lambda e^{-\lambda t} \tag{S1}$$



where $\lambda$ represents the rate parameter (average number of binding events per unit time).

- 2nd binding (3 unbound sites remaining): $\lambda_2 = 3k_{\text{on}}[L]$,
- 3rd binding (2 unbound sites remaining): $\lambda_3 = 2k_{\text{on}}[L]$,
- 4th binding (1 unbound site remaining): $\lambda_4 = k_{\text{on}}[L]$

#### S1.2 Probability Calculation of Binding Events

At most two binding events are observed in a single simulation (duration $T$). Thus, we calculate probabilities for three mutually exclusive outcomes: no binding events ($P(0)$), one binding event ($P(1)$), and two binding events ($P(2)$).

##### S1.2.1 Probability of No Binding Events ($P(0)$)

The probability of no binding events over the full simulation time $T$ is derived using the properties of the Poisson distribution. For no binding to occur, the first binding event (a prerequisite for all subsequent bindings) must not take place during $T$, leading to:

$$P(0) = e^{-4k_{\text{on}}[L]T} \tag{S2}$$

##### S1.2.2 Probability of One Binding Event ($P(1)$)

A single binding event requires two concurrent conditions:

- 1. The first binding occurs at time $t_1$ (where $0 < t_1 < T$), described by the exponential PDF for $\lambda_1$: $4k_{\text{on}}[L]e^{-4k_{\text{on}}[L]t_1}$,
- 2. No second binding event occurs in the remaining time $T - t_1$, with probability $e^{-\lambda_2(T-t_1)} = e^{-3k_{\text{on}}[L](T-t_1)}$.

Combining these independent conditions, the probability of one binding event is:

$$P(1) = 4k_{\text{on}}[L]e^{-4k_{\text{on}}[L]t_1} \cdot e^{-3k_{\text{on}}[L](T-t_1)} \tag{S3}$$

##### S1.2.3 Probability of Two Binding Events ($P(2)$)

Two binding events require three concurrent conditions:

- 1. The first binding occurs at time $t_2$ (where $0 < t_2 < T$), described by $4k_{\text{on}}[L]e^{-4k_{\text{on}}[L]t_2}$,

- 2. The second binding occurs at time $t_2'$ (where $t_2 < t_2' < T$), with the time interval $t_2' - t_2$ following the exponential PDF for $\lambda_2$: $3k_{\text{on}}[L]e^{-3k_{\text{on}}[L](t_2'-t_2)}$,

- 3. No third binding event occurs in the remaining time $T - t_2'$, with probability $e^{-\lambda_3(T-t_2')} = e^{-2k_{\text{on}}[L](T-t_2')}$.

Combining these conditions, the probability of two binding events is:

$$P(2) = 4k_{\text{on}}[L]e^{-4k_{\text{on}}[L]t_2} \cdot 3k_{\text{on}}[L]e^{-3k_{\text{on}}[L](t_2'-t_2)} \cdot e^{-2k_{\text{on}}[L](T-t_2')} \tag{S4}$$

#### S1.3 Construction of the Log-Likelihood Function

The likelihood function quantifies the probability of observing the full set of simulation outcomes given $k_{\text{on}}$. For independent simulations, the likelihood function is the product of individual outcome probabilities; taking the natural logarithm (log-likelihood) converts products to sums, simplifying maximization while preserving monotonicity.

##### S1.3.1 Observed Simulation Outcomes

Define the following counts based on simulation data:

- $n_0$: Number of simulations with no binding events,

- $n_1$: Number of simulations with one binding event (binding times denoted $t_1^{(j)}$ for $j = 1, 2, ..., n_1$),

- $n_2$: Number of simulations with two binding events (binding times denoted $t_2^{(j)}$ and $t_2'^{(j)}$ for $j = 1, 2, ..., n_2$).

##### S1.3.2 Log-Likelihood Expansion

Substitute $P(0)$, $P(1)$, and $P(2)$ into the likelihood function and take the natural logarithm:

$$
\begin{aligned}
\ln \mathcal{L}(k_{\text{on}}) &= n_0 \ln P(0) + \sum_{j=1}^{n_1} \ln P(1) + \sum_{j=1}^{n_2} \ln P(2) \\
&= -4n_0 k_{\text{on}}[L]T + \sum_{j=1}^{n_1} \left[ \ln(4k_{\text{on}}[L]) - 4k_{\text{on}}[L]t_1^{(j)} - 3k_{\text{on}}[L](T - t_1^{(j)}) \right] \\
&\quad + \sum_{j=1}^{n_2} \left[ \ln(4k_{\text{on}}[L]) + \ln(3k_{\text{on}}[L]) - 4k_{\text{on}}[L]t_2^{(j)} - 3k_{\text{on}}[L](t_2'^{(j)} - t_2^{(j)}) - 2k_{\text{on}}[L](T - t_2'^{(j)}) \right]
\end{aligned}
\tag{S5}
$$

#### S1.4 MLE of the Association Rate Constant $k_{\text{on}}$

The MLE of $k_{\text{on}}$ is obtained by maximizing $\ln \mathcal{L}(k_{\text{on}})$, which requires setting the first derivative of the log-likelihood function with respect to $k_{\text{on}}$ to zero (the second derivative is negative, confirming a maximum).

##### S1.4.1 First Derivative of the Log-Likelihood

Differentiate $\ln \mathcal{L}(k_{\text{on}})$ with respect to $k_{\text{on}}$ (treating $[L]$ and $T$ as constants):

$$
\begin{aligned}
\frac{d \ln \mathcal{L}}{dk_{\text{on}}} &= -4n_0[L]T - [L] \sum_{j=1}^{n_1} \left[ 4t_1^{(j)} + 3(T - t_1^{(j)}) \right] - [L] \sum_{j=1}^{n_2} \left[ 4t_2^{(j)} + 3(t_2'^{(j)} - t_2^{(j)}) + 2(T - t_2'^{(j)}) \right] \\
&\quad + \frac{n_1}{k_{\text{on}}} + \frac{2n_2}{k_{\text{on}}}
\end{aligned}
\tag{S6}
$$

##### S1.4.2 Solving for $k_{\text{on}}$

Set $\frac{d\ln\mathcal{L}}{dk_{\text{on}}} = 0$ and rearrange terms to isolate $k_{\text{on}}$:

$$
\begin{aligned}
k_{\text{on}} &= \frac{n_1 + 2n_2}{4[L]Tn_0 + \sum_{j=1}^{n_1}\left[4[L]t_1^{(j)} + 3[L](T - t_1^{(j)})\right] + \sum_{j=1}^{n_2}\left[4[L]t_2^{(j)} + 3[L](t_2'^{(j)} - t_2^{(j)}) + 2[L](T - t_2'^{(j)})\right]} \\
&= \frac{1}{[L]}\frac{n_1 + 2n_2}{4Tn_0 + \sum_{j=1}^{n_1}\left[4t_1^{(j)} + 3(T - t_1^{(j)})\right] + \sum_{j=1}^{n_2}\left[4t_2^{(j)} + 3(t_2'^{(j)} - t_2^{(j)}) + 2(T - t_2'^{(j)})\right]}
\end{aligned}
\tag{S7}
$$

- **Numerator**: Total number of binding events across all simulations ($1 \times n_1 + 2 \times n_2$),
- **Denominator**: Total "effective binding time" scaled by $[L]$, accounting for the number of unbound sites available during each simulation interval.

#### S1.5 Standard Deviation of $k_{\text{on}}$ ($\sigma_{k_{\text{on}}}$)

The standard deviation of $k_{\text{on}}$ is estimated using the error propagation formula, based on the Fisher Information Matrix (a standard MLE uncertainty quantification method).

##### S1.5.1 Fisher Information Calculation

The Fisher Information $\mathcal{I}$ is the negative expected value of the second derivative of the log-likelihood. For observed counts $n_0, n_1, n_2$, the expected value equals the observed value:

$$
\mathcal{I} = -\mathbb{E}\left[\frac{d^2\ln\mathcal{L}}{dk_{\text{on}}^2}\right] = \frac{n_1 + 2n_2}{k_{\text{on}}^2} \tag{S8}
$$

##### S1.5.2 Standard Deviation Derivation

The variance of $k_{\mathrm{on}}$ is $\mathrm{Var}(k_{\mathrm{on}}) = 1/\mathcal{I}$. Taking the square root and substituting $k_{\mathrm{on}}$ from Eq. (S7) gives:

$$\begin{aligned}\sigma_{k_{\mathrm{on}}} &= \frac{\sqrt{n_1 + 2n_2}}{4[L]Tn_0 + \sum_{j=1}^{n_1}\left[4[L]t_1^{(j)} + 3[L](T - t_1^{(j)})\right] + \sum_{j=1}^{n_2}\left[4[L]t_2^{(j)} + 3[L](t_2'^{(j)} - t_2^{(j)}) + 2[L](T - t_2'^{(j)})\right]} \\ &= \frac{1}{[L]}\frac{\sqrt{n_1 + 2n_2}}{4Tn_0 + \sum_{j=1}^{n_1}\left[4t_1^{(j)} + 3(T - t_1^{(j)})\right] + \sum_{j=1}^{n_2}\left[4t_2^{(j)} + 3(t_2'^{(j)} - t_2^{(j)}) + 2(T - t_2'^{(j)})\right]}\end{aligned} \tag{S9}$$

### S2 A baseline study using an alanine dipeptide (Ala2) as model system for potential impact of HMR on system dynamics

#### S2.1 Basic experimental design

For two schemes, we performed two independent 1 $\mu$s simulations: (1) using the CHARMM36m force field with HMR, at a time step of 2 fs; and (2) using the CHARMM36m force field without HMR, also at a time step of 2 fs. We considered the distances between all heavy atoms as features. To ensure consistency of comparison, we merged all four feature trajectories and performed dimensionality reduction using TICA and subsequent k-means clustering. We then constructed Markov state models (MSMs) and the Robust Perron-cluster cluster analysis (PCCA+)· for each scheme.

#### S2.2 Detailed Parameters

We obtained a total of 45 distance pairs between heavy atoms. We set the feature extraction frequency to 1 ps/frame, the TICA lag time to 1 ps, k-means to cluster the initial trajectories into 100 classes on the first 10 dimensions of TIC, the MSM lag time to 5 ps, and the macrostate classes to 5. We calculated the probability of the macrostate and the mean first passage time (MFPT) between macrostates.

#### S2.3 Results

The results showed that the metastable state distributions and interstate transition rates were almost identical for schemes with and without HMR enabled (Fig. S27). This validation confirms that using HMR with a 2 fs time step does not alter the fundamental dynamics or free energy landscape of the peptide system.

### Supplementary Tables

**Table S1:** Summary of biological insights

| System | Binding mode | Binding Pathway and Rate | $\chi_1^{W158}$ Free Energy Landscape | Dissociation Pathway and Rate | Absolute Binding Free Energy |
|---|---|---|---|---|---|
| MhOR5-EOL | Yes | Yes | Yes | Yes | Yes |
| MhOR5-DEET | Yes | Yes | Yes | Yes | Yes |
| MhOR5$_{apo}$ | - | - | Yes | - | - |

**Table S2:** Summary of all simulations timescales, replicas and motivation

| No. | Algorithm | Replicas | Windows | Timescales for each replica (window) (ns) | System | Motivation |
|---|---|---|---|---|---|---|
| 1 | Unbiased | 3 | 0 | 1000 | MhOR5-EOL | Binding mode |
| | | 3 | 0 | 1000 | MhOR5-DEET | |
| 2 | | 10 | 0 | 1000 | MhOR5 and 10 EOL | Binding pathway and time |
| | | 10 | 0 | 1000 | MhOR5 and 10 DEET | |
| 3 | RAMD | 10 | 0 | up to 200 | MhOR5-EOL | Dissociation pathway |
| | | 10 | 0 | up to 200 | MhOR5-DEET | |
| 4 | ABMD | 10 | 0 | up to 200 | MhOR5-EOL | Dissociation pathway |
| | | 10 | 0 | up to 200 | MhOR5-DEET | |
| 5 | Adaptive Sampling | 24 | 0 | 50 | MhOR5-EOL | Slow variables |
| 6 | OPES | 1 | 0 | 1000 | MhOR5-EOL | $\chi_1^{W158}$ and $\chi_2^{W158}$ Free Energy Landscape |
| | | 1 | 0 | 1000 | MhOR5-DEET | |
| | | 1 | 0 | 1000 | MhOR5$_{apo}$ | |
| 7 | OPES-Flooding | 20 | 0 | up to 100 | MhOR5-EOL | Dissociation pathway and time |
| | | 20 | 0 | up to 400 | MhOR5-DEET | |
| 8 | FEP | 1 | 21 | 40-120 | MhOR5-EOL/EOL | ABFE |
| | | 1 | 21 | 20 | MhOR5-DEET/DEET | |

**Table S3:** Binding pathways with $k_{\text{on}}$ and $t_{\text{on}}$, and their associated uncertainties from maximum likelihood estimation

| Ligand | Pathway | No. of Events | $\tau_{on}^{MLE}$ ($\mu$s) | $k_{on}^{MLE}$ ($10^6 M^{-1} s^{-1}$) |
|---|---|---|---|---|
| EOL | From solvent | 1 | $4.94 \pm 3.49$ | $5.27 \pm 3.73$ |
| EOL | From membrane | 1 | | |
| DEET | From solvent | 2 | $1.61 \pm 0.66$ | $16.18 \pm 6.61$ |
| DEET | From membrane | 4 | | |

**Table S4:** Definition of candidate features involving distance terms entered into KTICA

| No. | Feature name | Definition |
|---|---|---|
| 1 | pocket | Y99,S151,G154,W155,W158,I213,Y380,Y383 |
| 2 | d0 | Distance between center of ligand and center of Y91 |
| 3 | d1 | Distance between center of ligand and center of S151 |
| 4 | d2 | Distance between center of ligand and center of G154 |
| 5 | d3 | Distance between center of ligand and center of W155 |
| 6 | d4 | Distance between center of ligand and center of W158 |
| 7 | d5 | Distance between center of ligand and center of I213 |
| 8 | d6 | Distance between center of ligand and center of Y380 |
| 9 | d7 | Distance between center of ligand and center of Y383 |
| 10 | dl-p | Distance between center of ligand and center of pocket |
| 11 | $e^{-d*}$ | Negative exponent power of d*, d* represents the above distance |

**Table S5:** ABFE calculations of EOL and DEET using FEP. $\Delta G$ unit in kcal/mol

| System | $\Delta G_1$ | $\Delta G_3$ | $\Delta G_4$ | $\Delta G_{Binding}$ | $\Delta\Delta G_{\text{DEET - EOL}}$ |
|---|---|---|---|---|---|
| MhOR5 - EOL | 20.24 ± 0.17 | 9.27 | 2.58 ± 0.14 | -8.39 ± 0.22 | -2.86 ± 0.27 |
| MhOR5 - DEET | 28.16 ± 0.083 | 9.73 | 7.18 ± 0.16 | -11.25 ± 0.16 | |

### Supplementary Figures

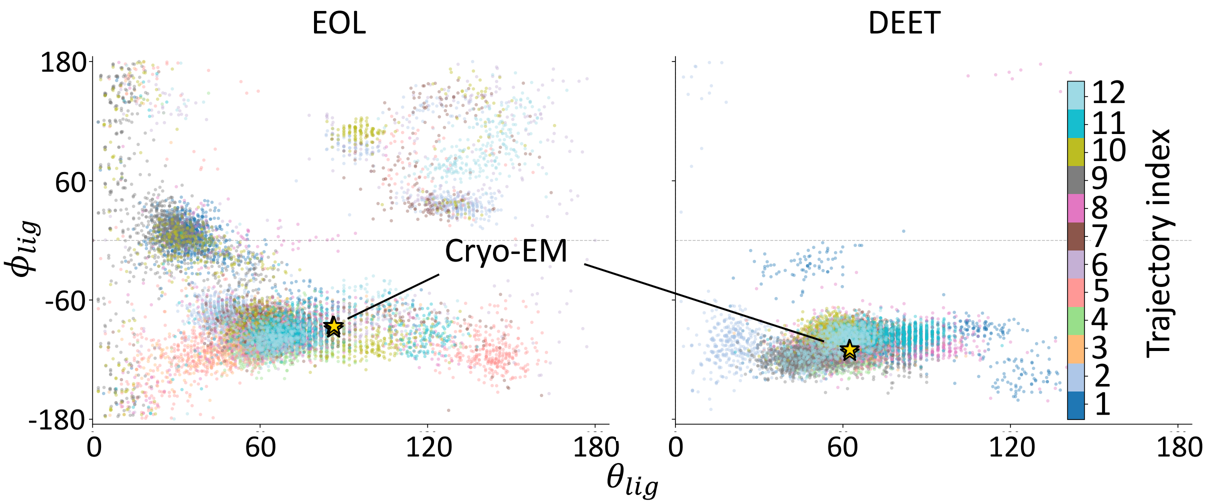


**Figure S1:** The trajectories of the $\theta_{lig}$ and $\phi_{lig}$ of ligand of independent replicas. The projection positions of cryo-EM structure are shown in yellow star.

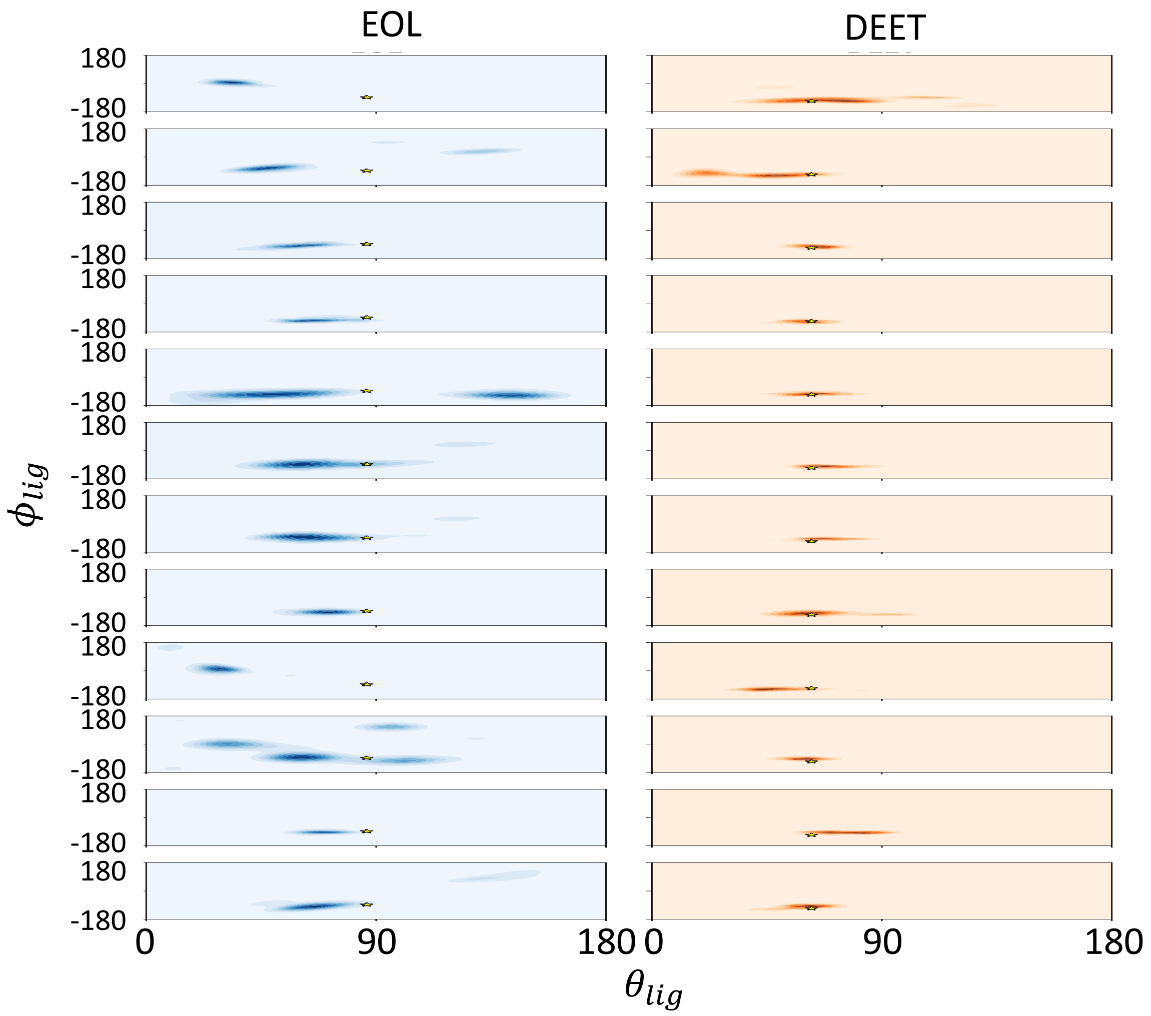


**Figure S2:** The distributions of the $\theta_{lig}$ and $\phi_{lig}$ of ligand of independent replicas. The projection positions of cryo-EM structure are shown in yellow star.

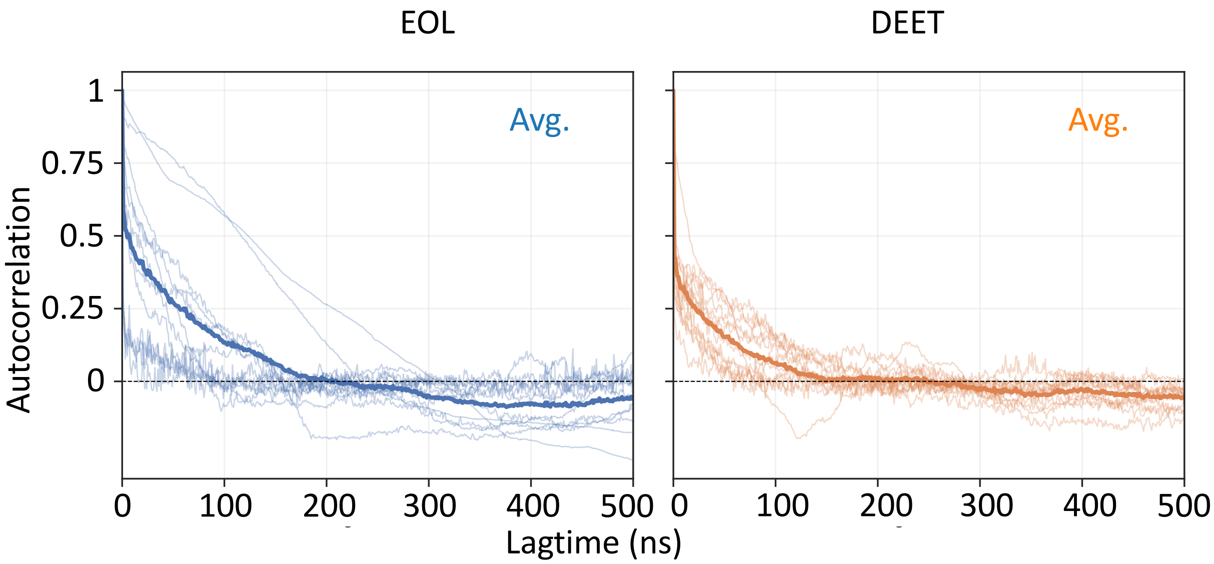


**Figure S3:** The autocorrelation plots of both $\theta_{lig}$ and $\phi_{lig}$ of EOL and DEET.

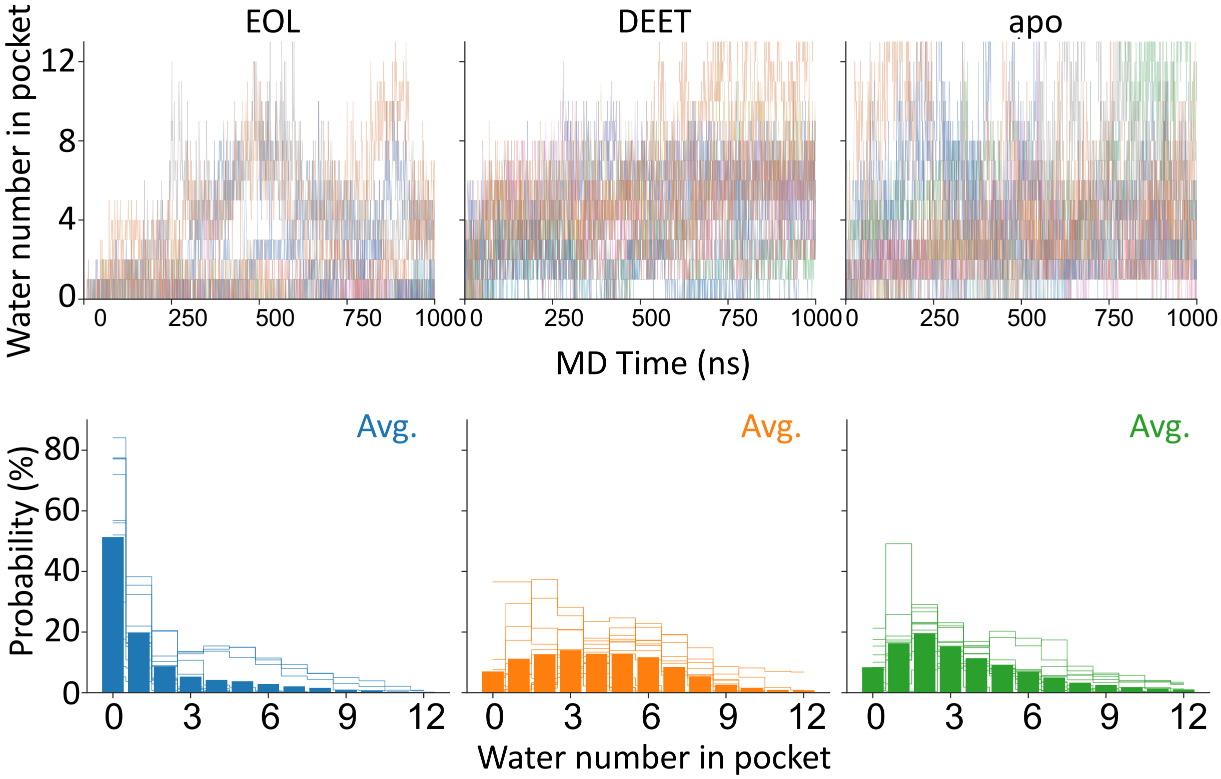


**Figure S4:** The trajectories and histograms of water number in pocket of independent replicas. In the histogram plots, the transparent bars represent the histograms of independent replicas and the solid bar represents the average histogram.

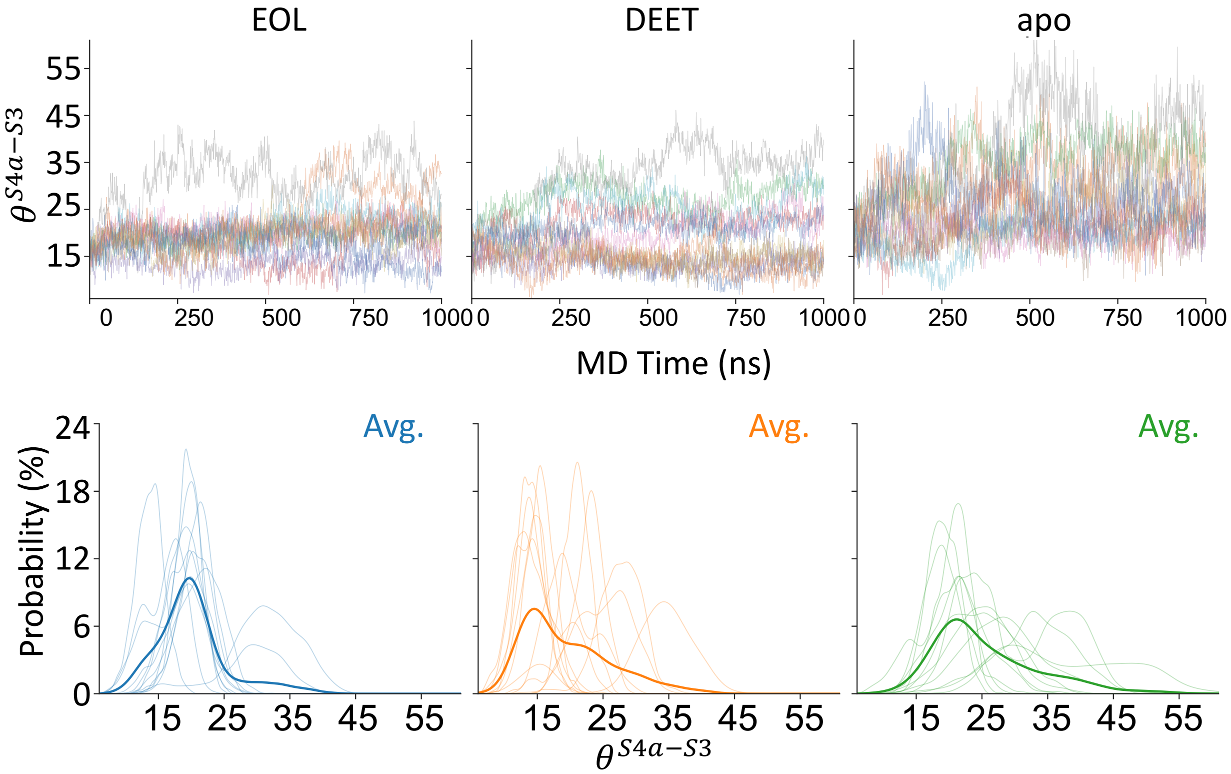


**Figure S5:** The trajectories and distributions of $\theta^{S4a-s3}$ of independent replicas. In the distribution plots, the transparent lines represent the distributions of independent replicas and the solid line represents the average distribution.

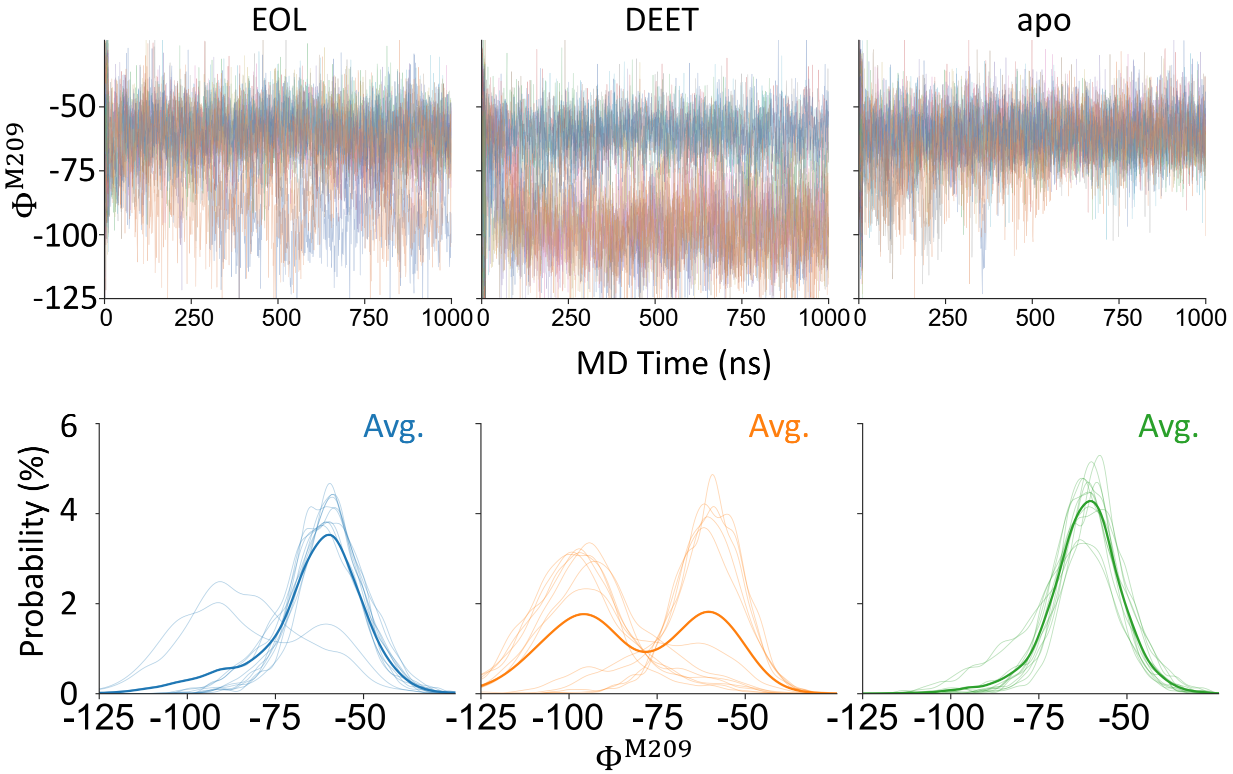


**Figure S6:** The trajectories and distributions of $\Phi^{M209}$ of independent replicas. In the distribution plots, the transparent lines represent the distributions of independent replicas and the solid line represents the average distribution.

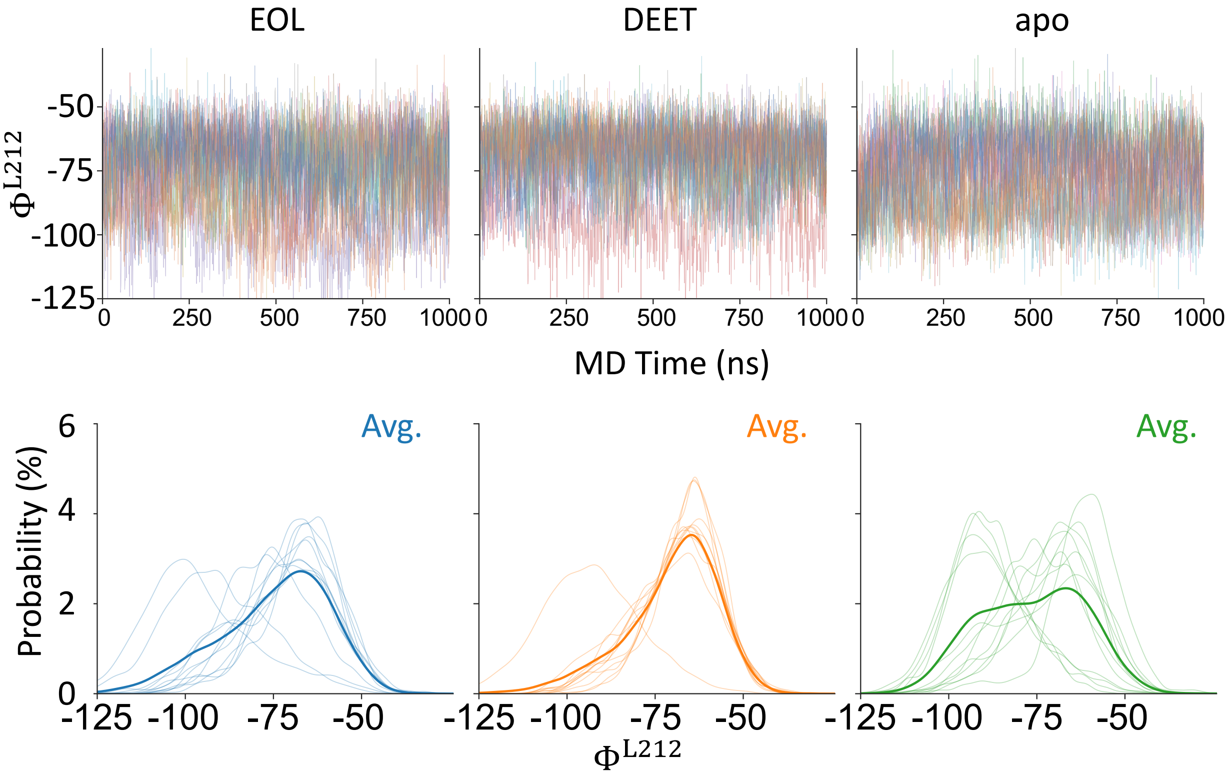


**Figure S7:** The trajectories and distributions of $\Phi^{L212}$ of independent replicas. In the distribution plots, the transparent lines represent the distributions of independent replicas and the solid line represents the average distribution.

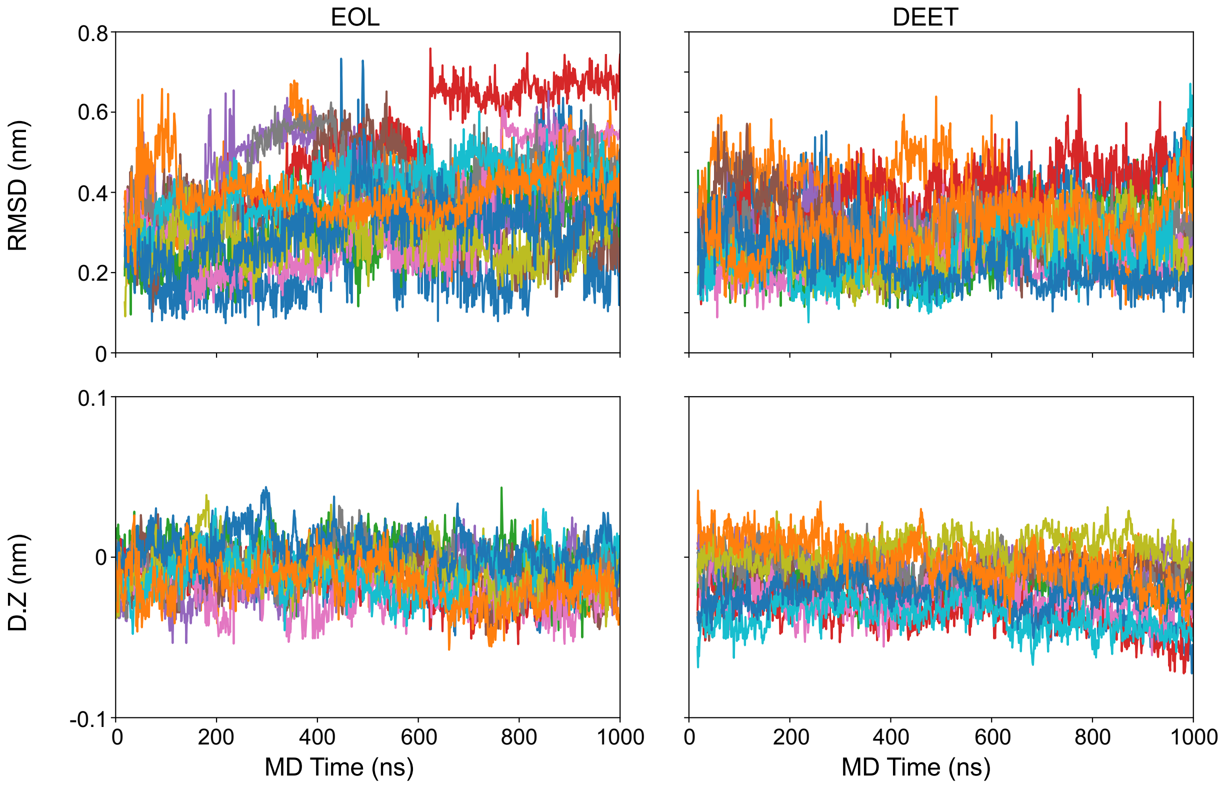


**Figure S8:** RMSD and vertical distances from membrane center of EOL and DEET in unbiased MD.

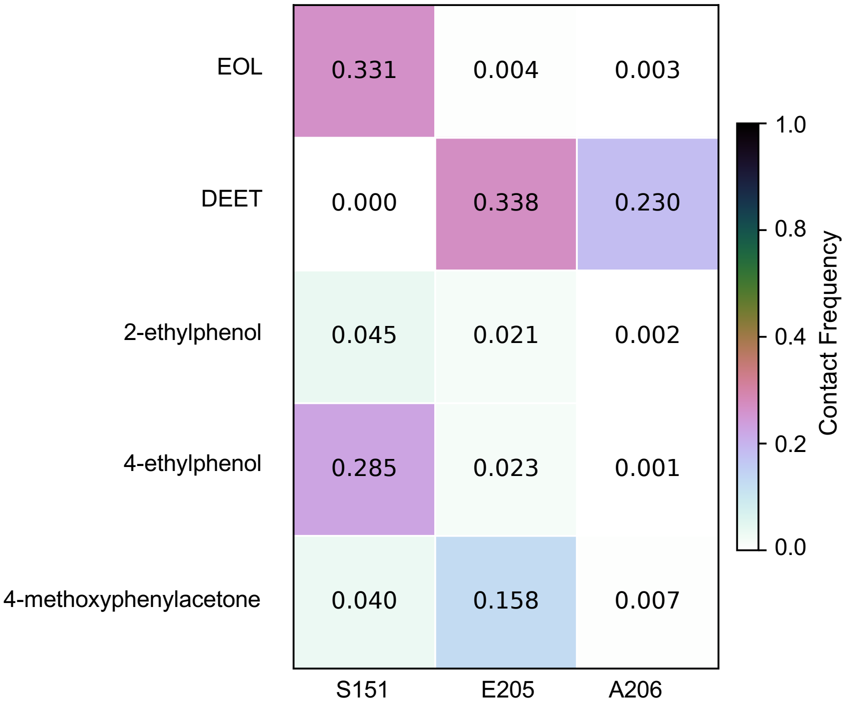


**Figure S9:** Frequencies of direct and water-mediated hydrogen bonds formed by five ligands during unbiased MD simulations. The S151 column reports direct ligand–S151 hydrogen bonds, whereas the E205 and A206 columns report water-mediated ligand–water–residue hydrogen bonds. Cell values indicate the fraction of trajectory frames containing the corresponding interaction.

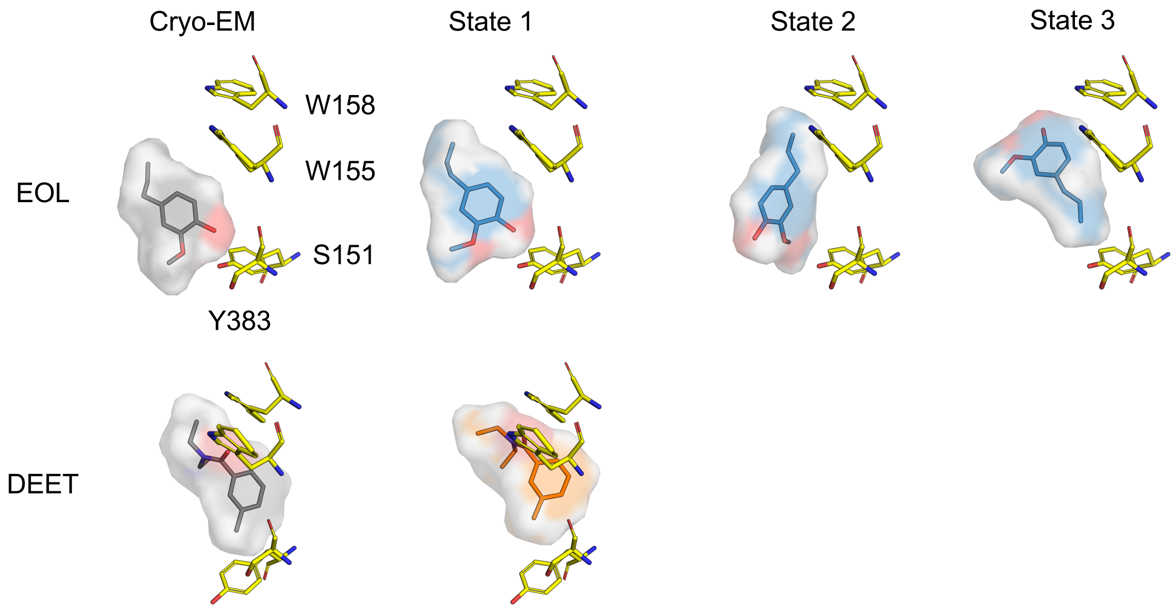


**Figure S10:** The main conformation from unbiased MD simulations compared with the cryo-EM structures.

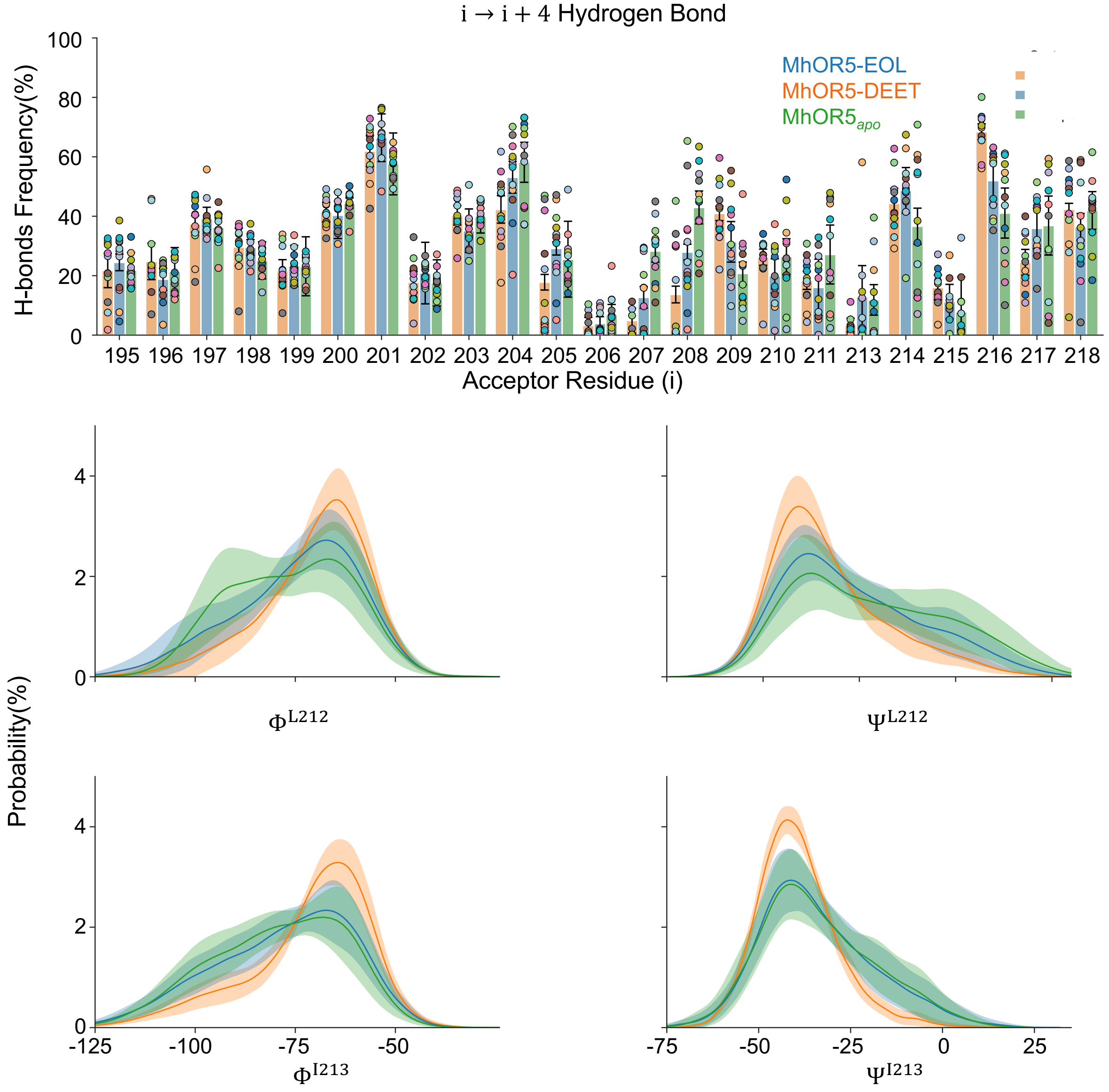


**Figure S11:** P216 breaks the formation of hydrogen bonds, thereby affecting the $\Phi/\Psi$ of the hydrogen bond acceptor residues. The twelve independent trajectories are marked with scatter points of distinct colors, and the 95% CI for each trajectory is calculated to serve as error bars.

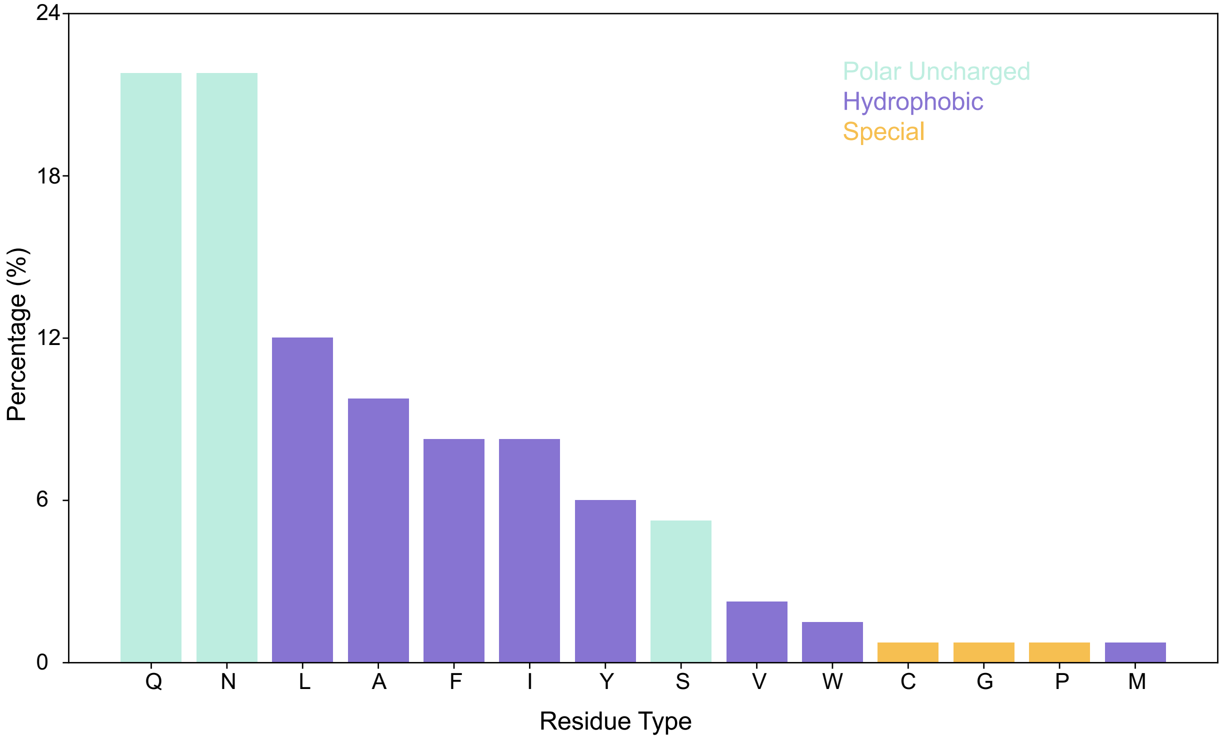


**Figure S12:** Multiple sequence alignment (MSA) of the S4 region was performed across 134 olfactory receptor sequences to characterize the identity and physicochemical properties of residues corresponding to P216.

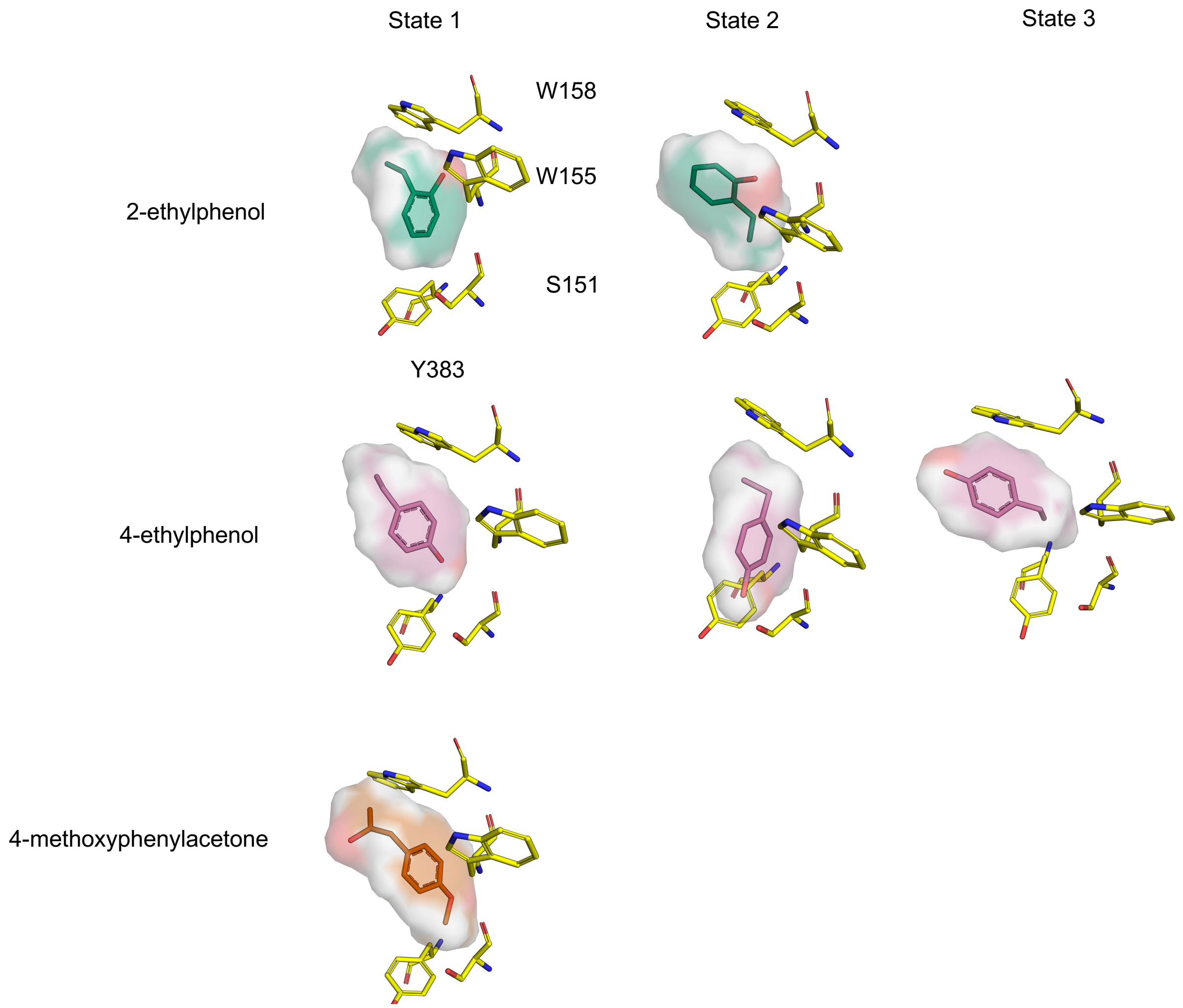


**Figure S13:** Representative binding poses of 2-ethylphenol, 4-ethylphenol, and 4-methoxyphenylacetone for the major orientational states identified from the two-dimensional distributions of $\theta_{lig}$ and $\phi_{lig}$. Ligands are shown as colored sticks and surfaces, and surrounding binding-pocket residues are shown as yellow sticks.

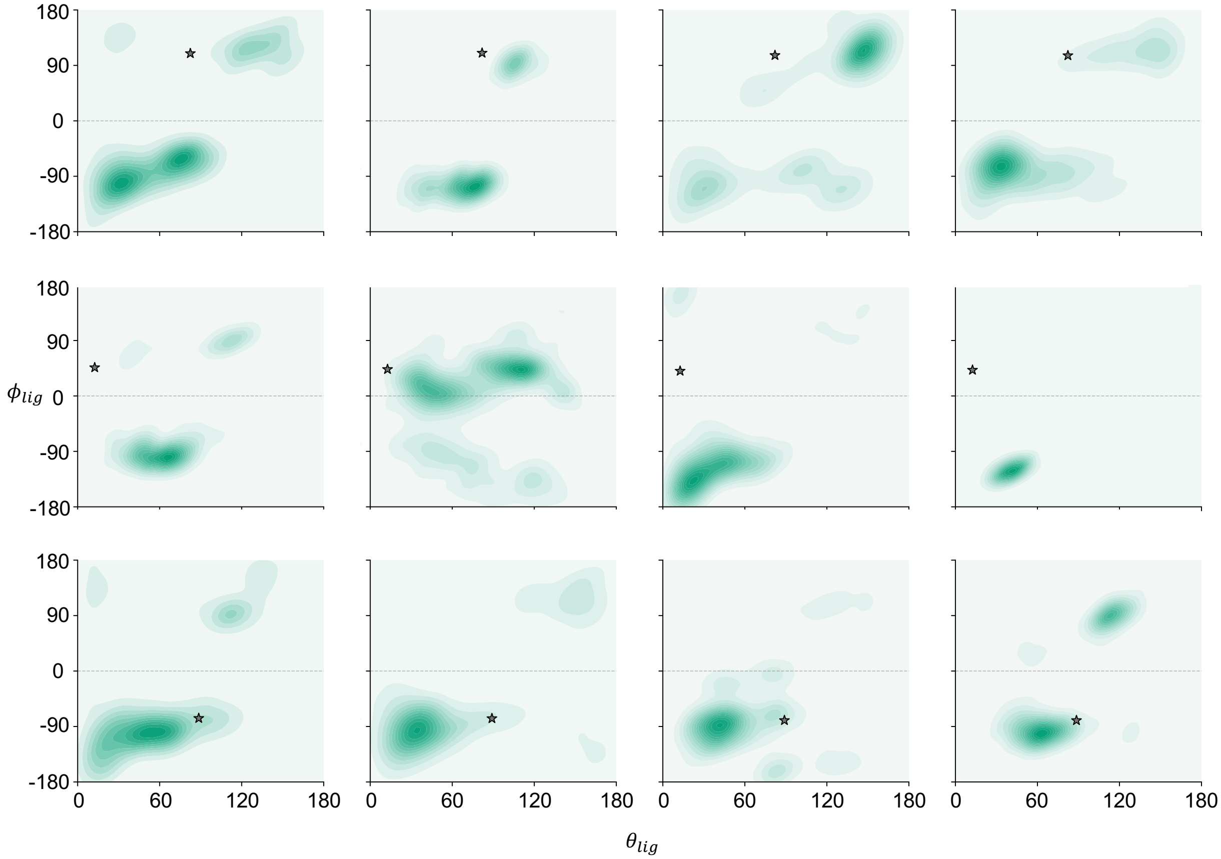


**Figure S14:** Two-dimensional distributions of $\theta_{lig}$ and $\phi_{lig}$ for 2-ethylphenol in the 12 ligand–subunit trajectories. Stars indicate the corresponding initial poses selected from AlphaFold 3 predictions.

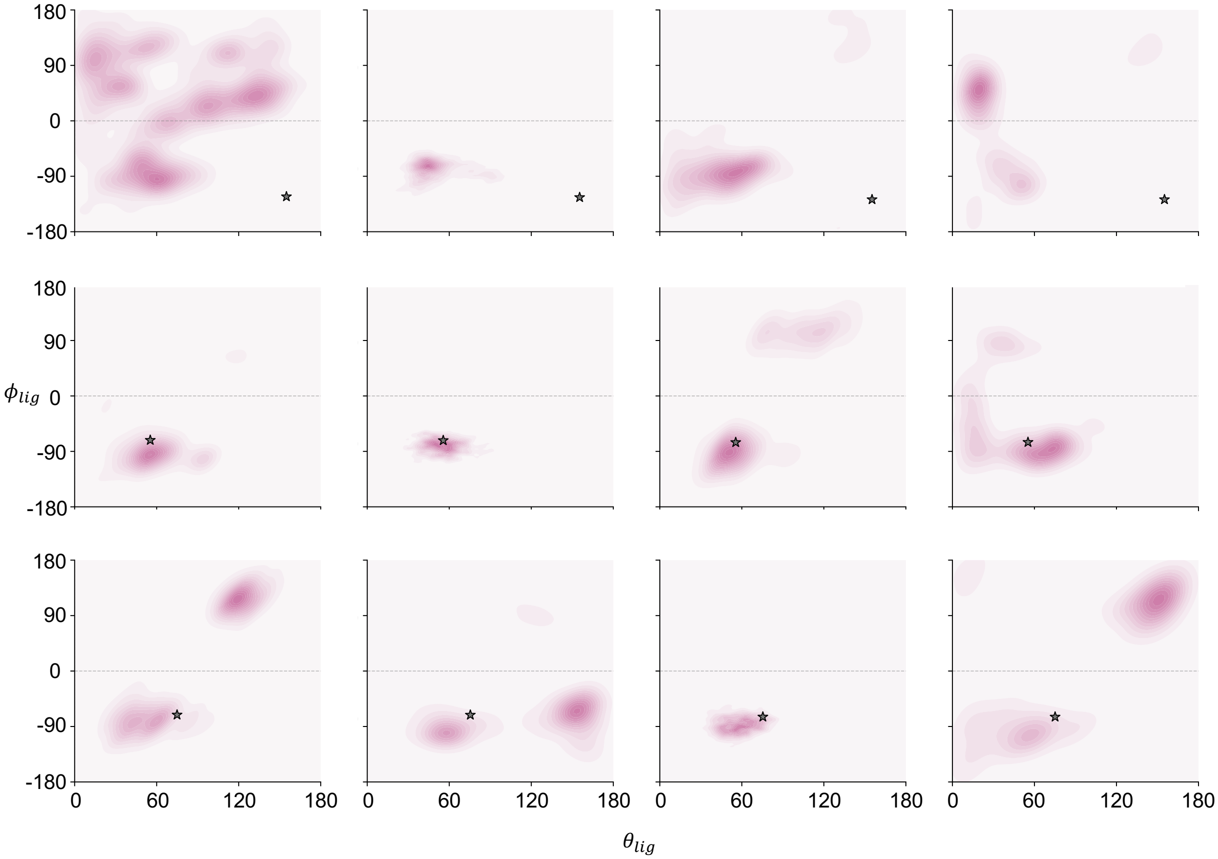


**Figure S15:** Two-dimensional distributions of $\theta_{lig}$ and $\phi_{lig}$ for 4-ethylphenol in the 12 ligand–subunit trajectories. Stars indicate the corresponding initial poses selected from AlphaFold 3 predictions.

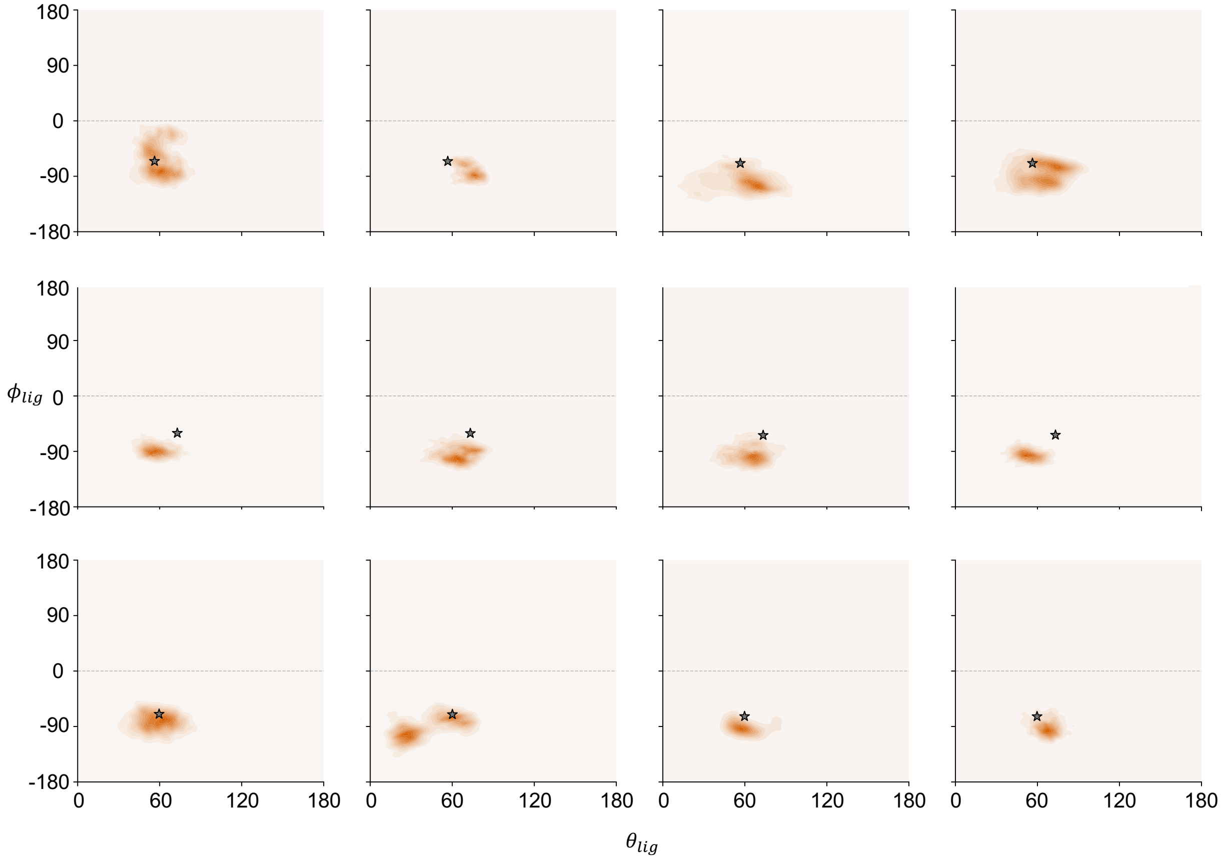


**Figure S16:** Two-dimensional distributions of $\theta_{lig}$ and $\phi_{lig}$ for 4-methoxyphenylacetone in the 12 ligand–subunit trajectories. Stars indicate the corresponding initial poses selected from AlphaFold 3 predictions.

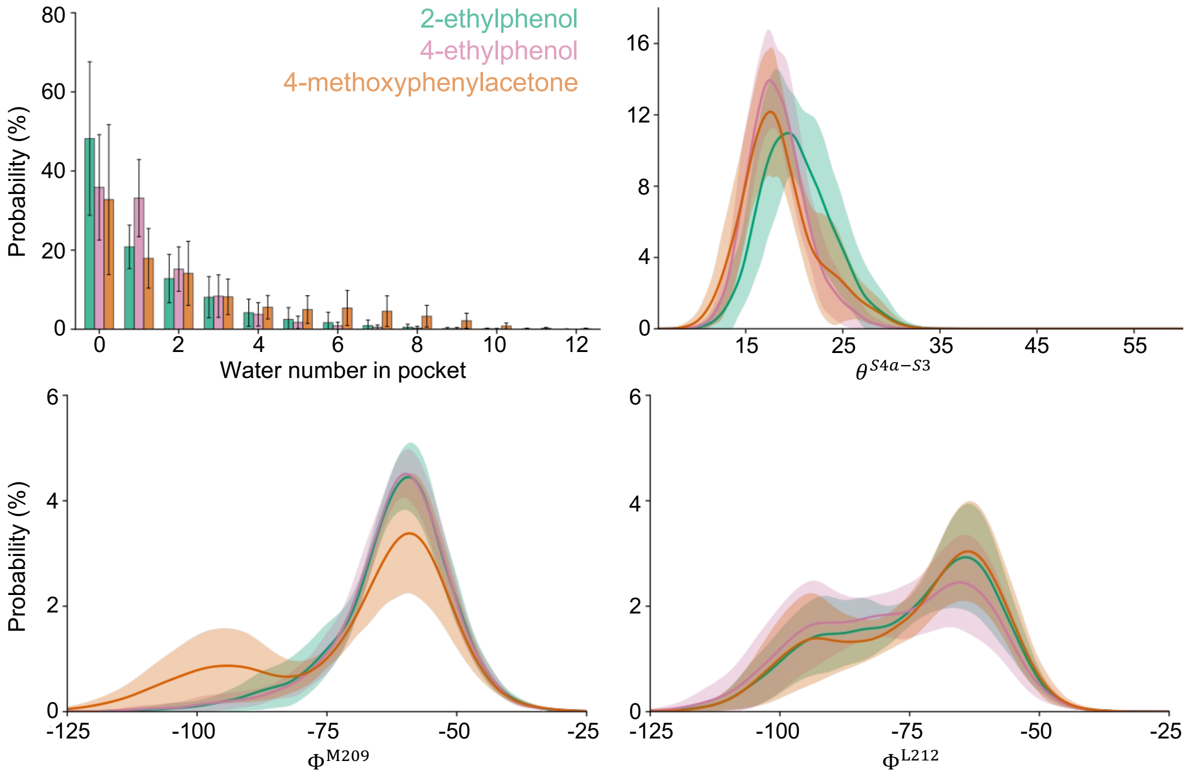


**Figure S17:** Binding-pocket hydration and S4 conformational dynamics for 2-ethylphenol, 4-ethylphenol, and 4-methoxyphenylacetone. The panels show the pocket water-number distributions, the probability distributions of the S4a–S3 angle ($\theta^{S4a-S3}$), and the backbone $\Phi$-angle distributions of M209 and L212. Lines or bars show the mean distributions across the 12 ligand–subunit trajectories, and shaded regions or error bars denote trajectory-based confidence intervals.

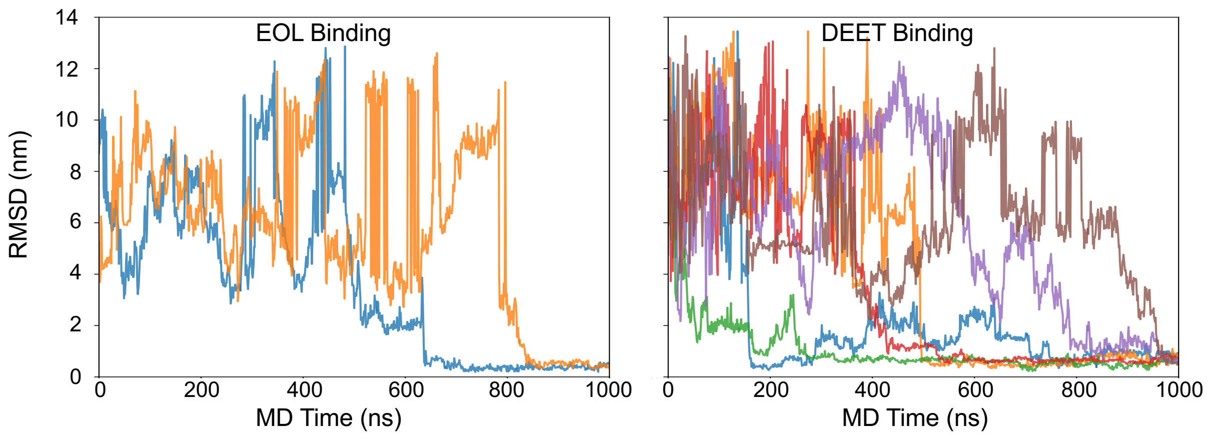


**Figure S18:** RMSD of EOL (left) and DEET (right) in the successful spontaneous binding trajectories compared with the cryo-EM structure, respectively. The accidental jumps in RMSD are not non-physical, but rather caused by PBC.

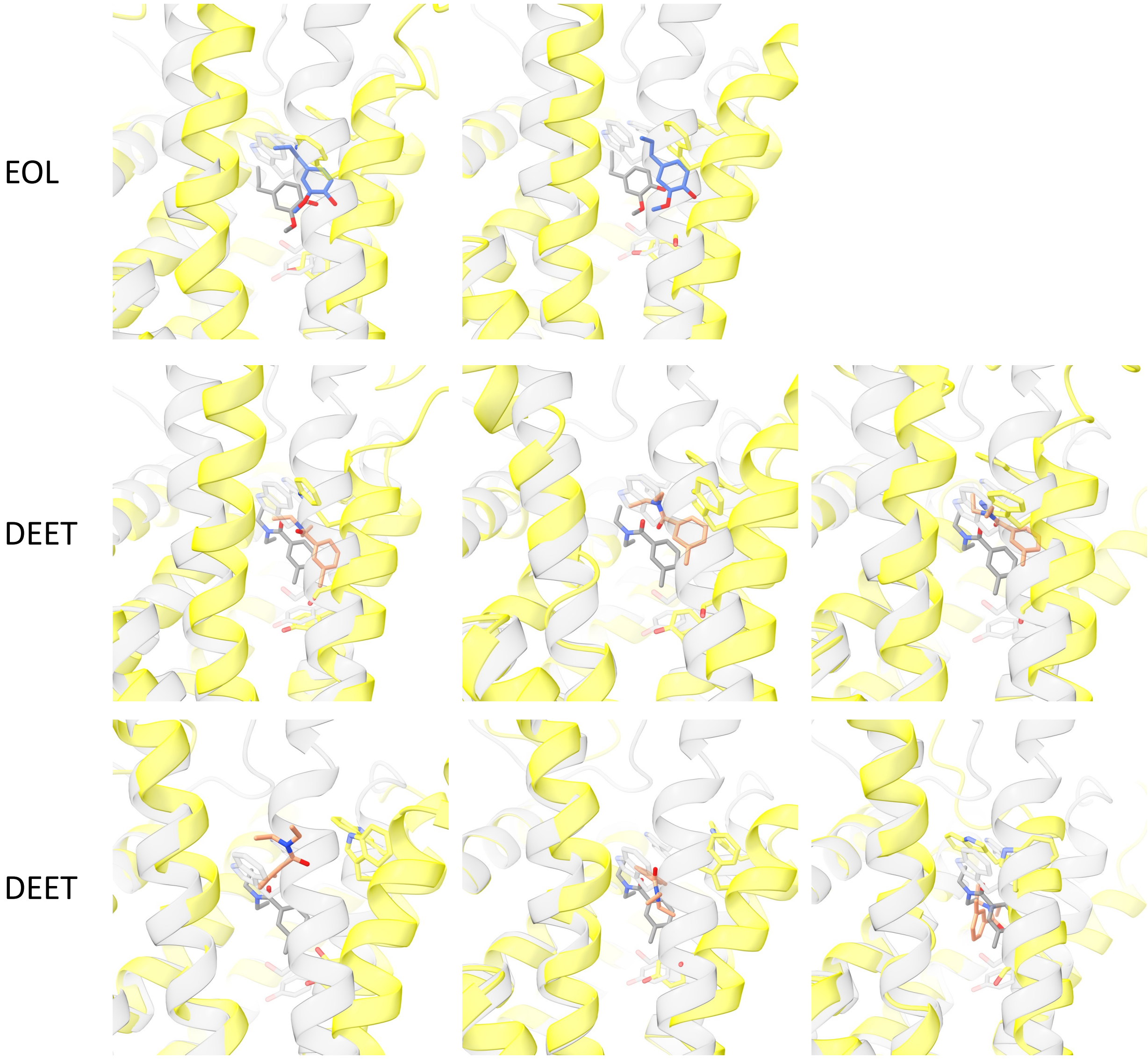


**Figure S19:** Binding poses of EOL and DEET in the spontaneous binding trajectories compared with the cryo-EM structures.

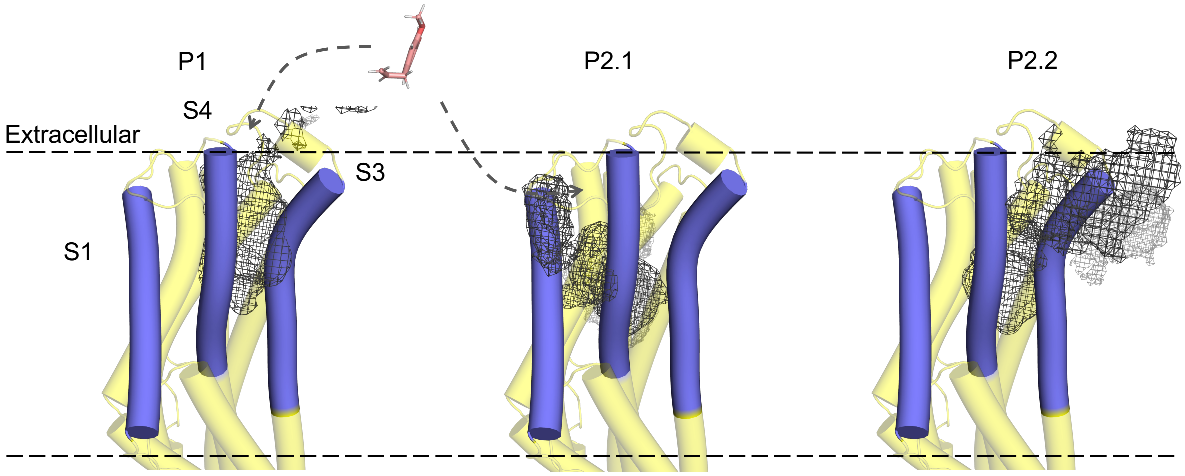


**Figure S20:** Both ligand binding and unbinding processes have three detailed pathways.

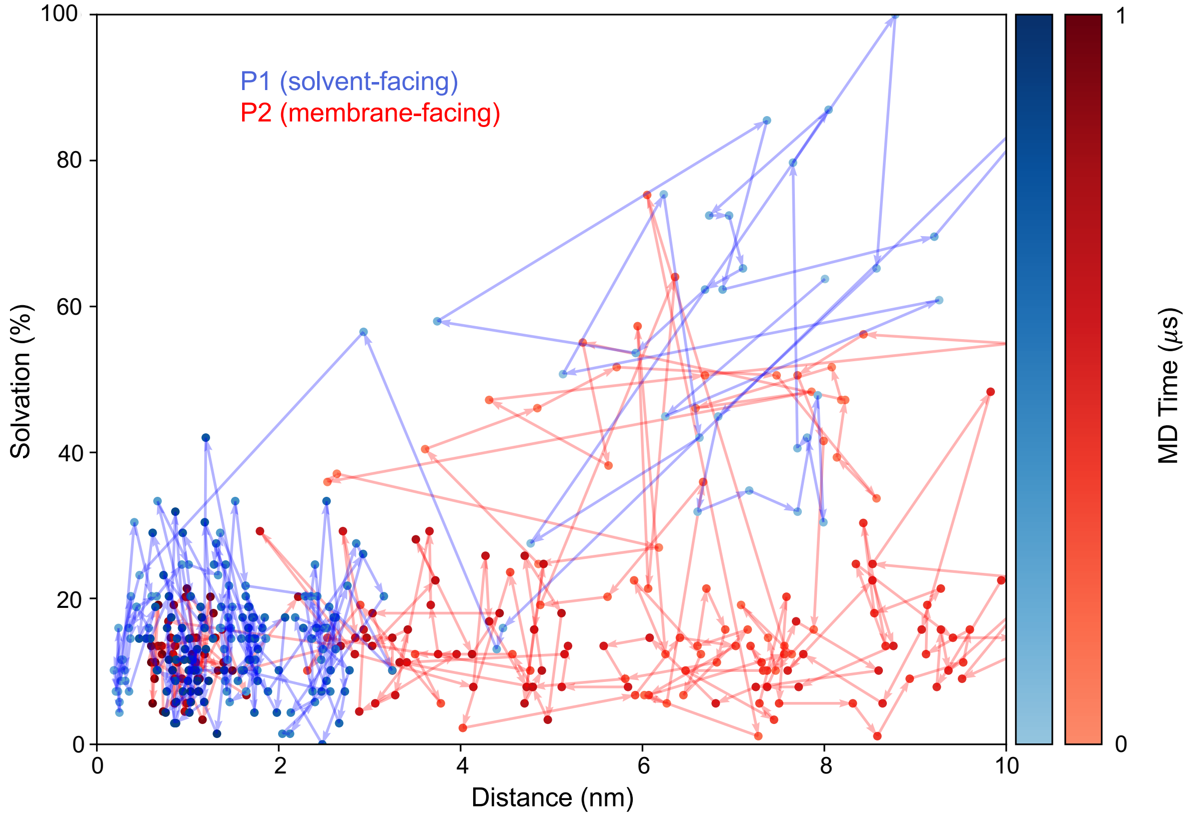


**Figure S21:** One representative trajectory for ligand binding of two parameters via pathways P1 and P2: the distance between the ligand center and the binding cavity center, and ligand solvent accessibility. The blue line denotes the solvent-facing pathway (P1), while the red line denotes the membrane-facing pathway (P2).

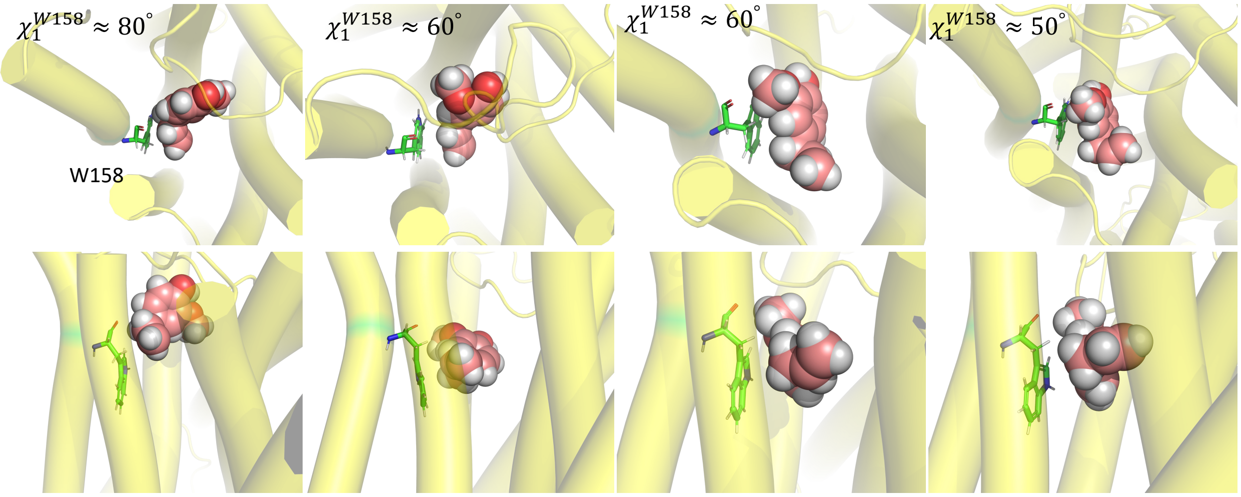


**Figure S22:** Snapshots of four W158 crossing events. Top (upper) and side (lower) views are shown. The protein is rendered as cartoons, the ligand as light-pink spheres, and W158 as green sticks.

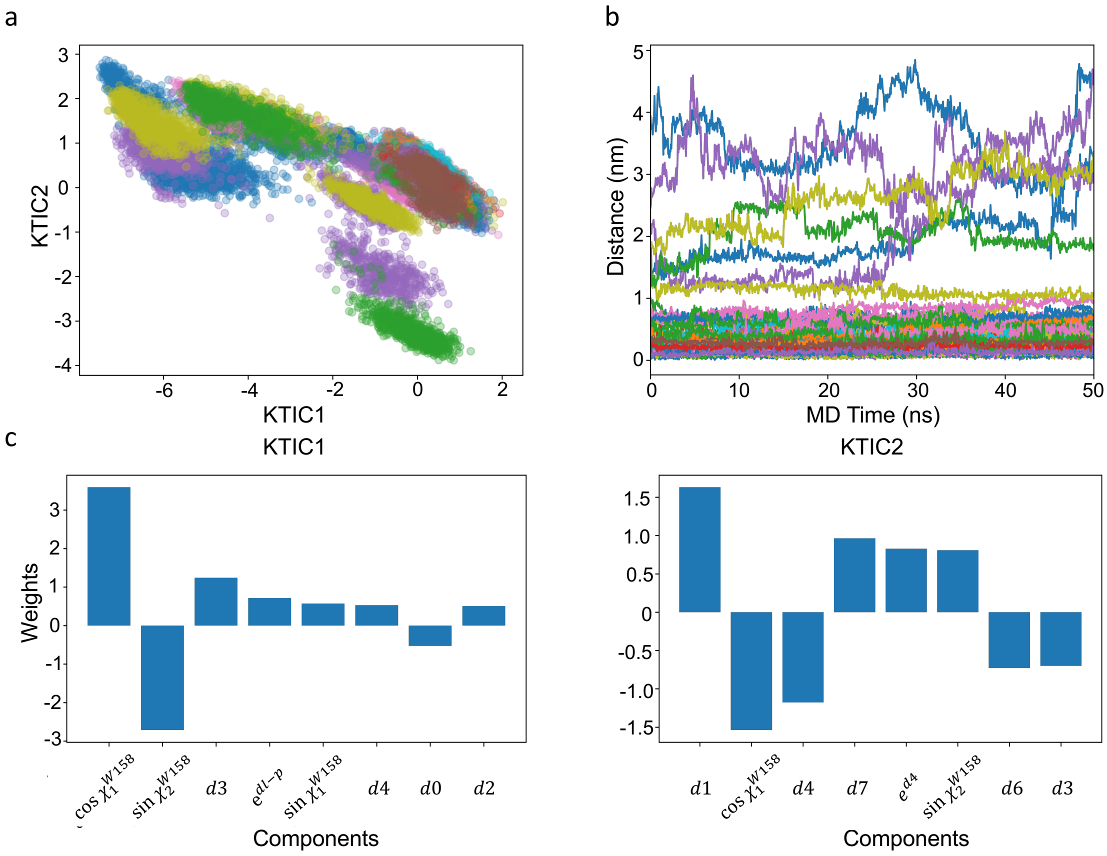


**Figure S23:** KTICA analysis of 24 adaptive sampling MD trajectories. (a,b) Projections of the 24 adaptive sampling trajectories onto the two leading KTICA components (KTIC1 and KTIC2) and onto the $d_{l-p}$ coordinate, respectively. (c) The first eight KTICA components and their corresponding weights contributing to KTIC1 and KTIC2.

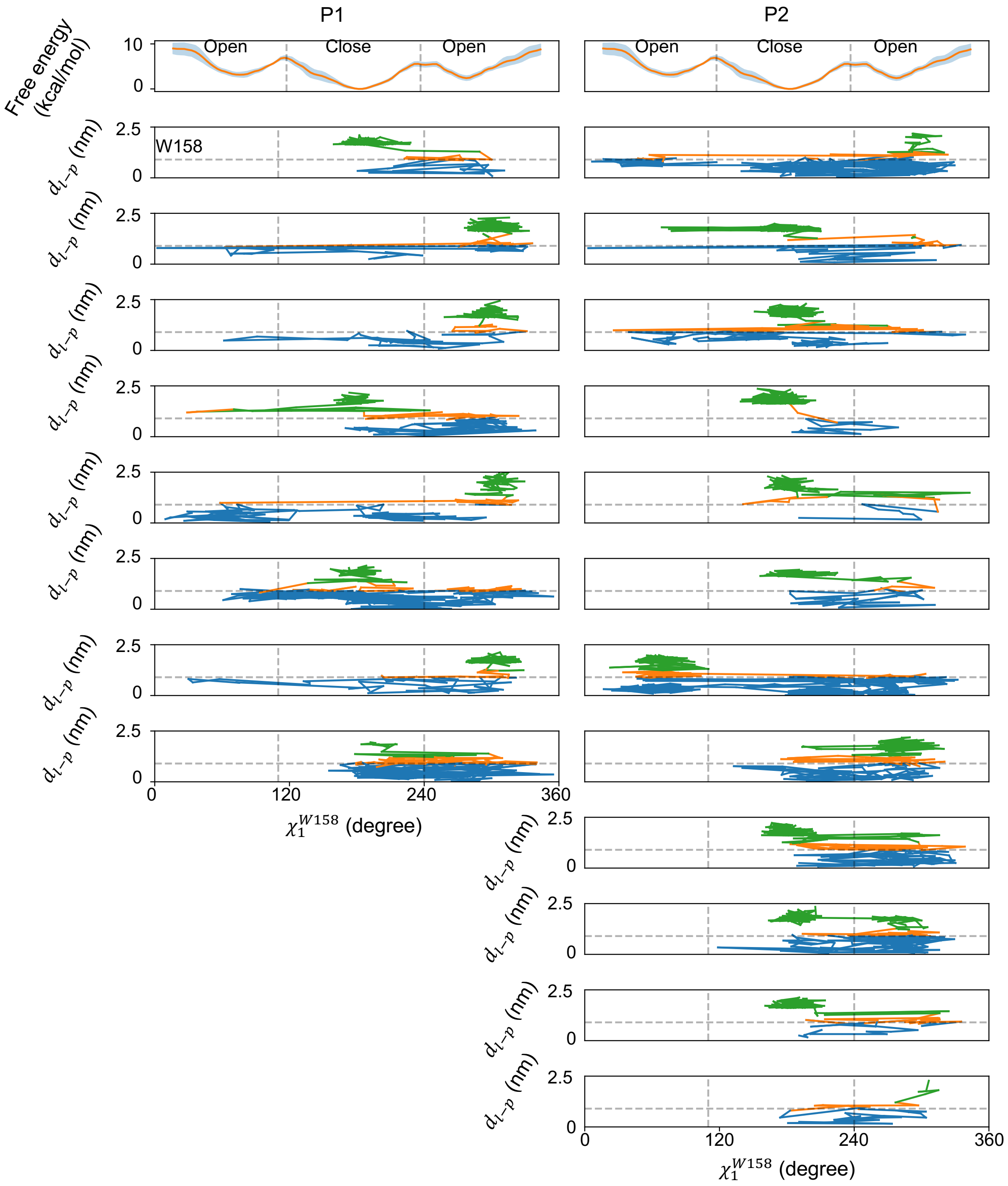


**Figure S24:** Projection of EOL dissociation trajectories onto $\chi_1^{W158}$ and $d_{l-p}$ coordinates. Twenty OPES-Flooding dissociation trajectories are projected onto the $\chi_1^{W158}$ dihedral angle and the $d_{l-p}$ coordinate and compared with the corresponding $\chi_1^{W158}$ free energy landscape. Trajectory segments are colored blue when the ligand does not cross W158, orange during W158 crossing, and green after complete crossing of W158.

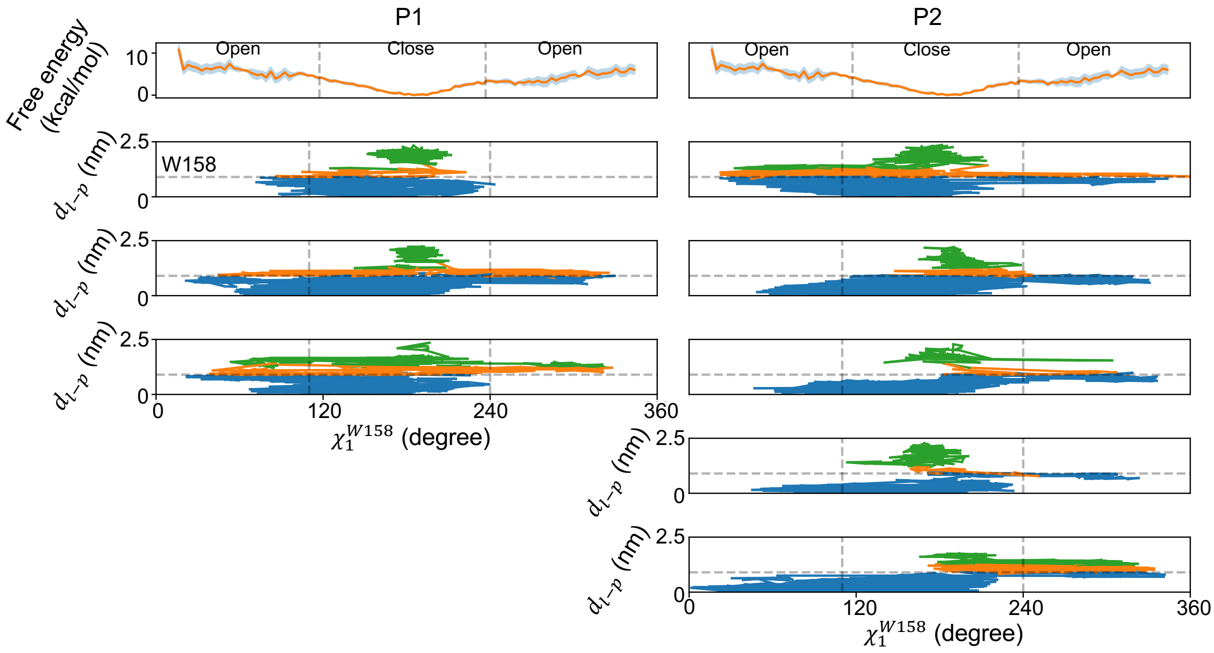


**Figure S25:** Projection of DEET dissociation trajectories onto $\chi_1^{W158}$ and $d_{l-p}$ coordinates. 8 OPES-Flooding dissociation trajectories are projected onto the $\chi_1^{W158}$ dihedral angle and the $d_{l-p}$ coordinate and compared with the corresponding $\chi_1^{W158}$ free energy landscape. Trajectory segments are colored blue when the ligand does not cross W158, orange during W158 crossing, and green after complete crossing of W158.

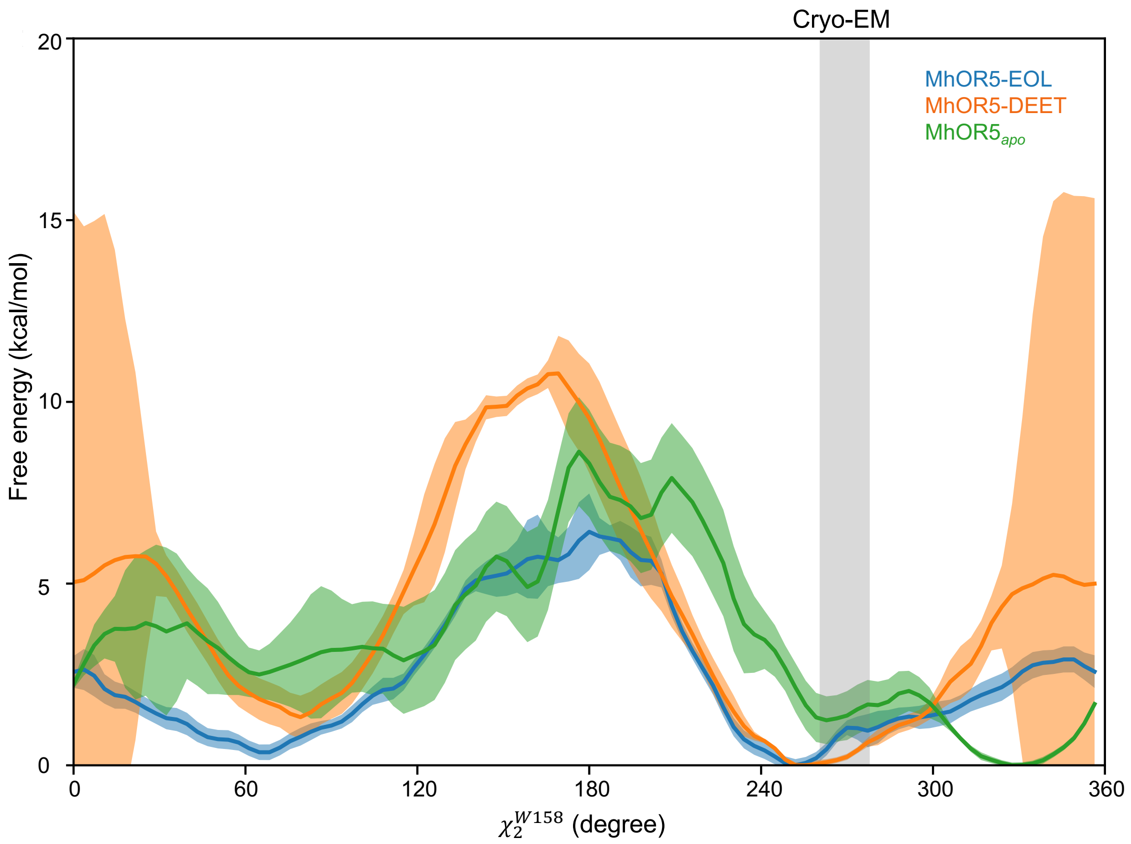


**Figure S26:** Free energy landscapes of $\chi_2^{W158}$ in MhOR5–EOL, MhOR5–DEET, and MhOR5$_{apo}$ systems. The uncertainties 95% CI were obtained using block analysis with 10 blocks; solid lines denote the mean free energy values and shaded regions indicate the associated uncertainties. The gray shaded area marks the range of $\chi_2^{W158}$ angles observed in three cryo-EM structures.

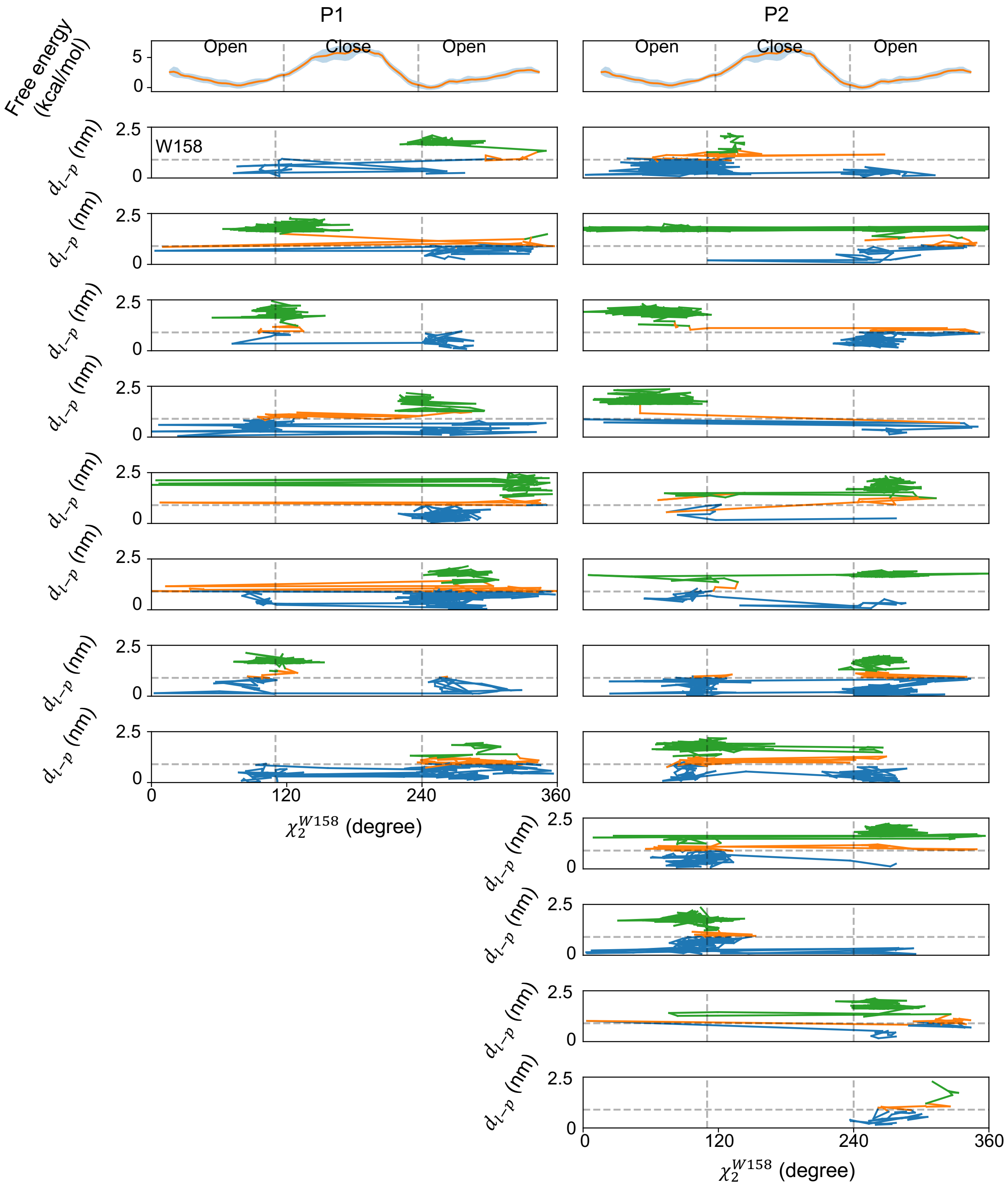


**Figure S27:** Projection of EOL dissociation trajectories onto $\chi_2^{W158}$ and $d_{l-p}$ coordinates. Twenty OPES-Flooding dissociation trajectories are projected onto the $\chi_2^{W158}$ dihedral angle and the $d_{l-p}$ coordinate and compared with the corresponding $\chi_2^{W158}$ free energy landscape. Trajectory segments are colored blue when the ligand does not cross W158, orange during W158 crossing, and green after complete crossing of W158.

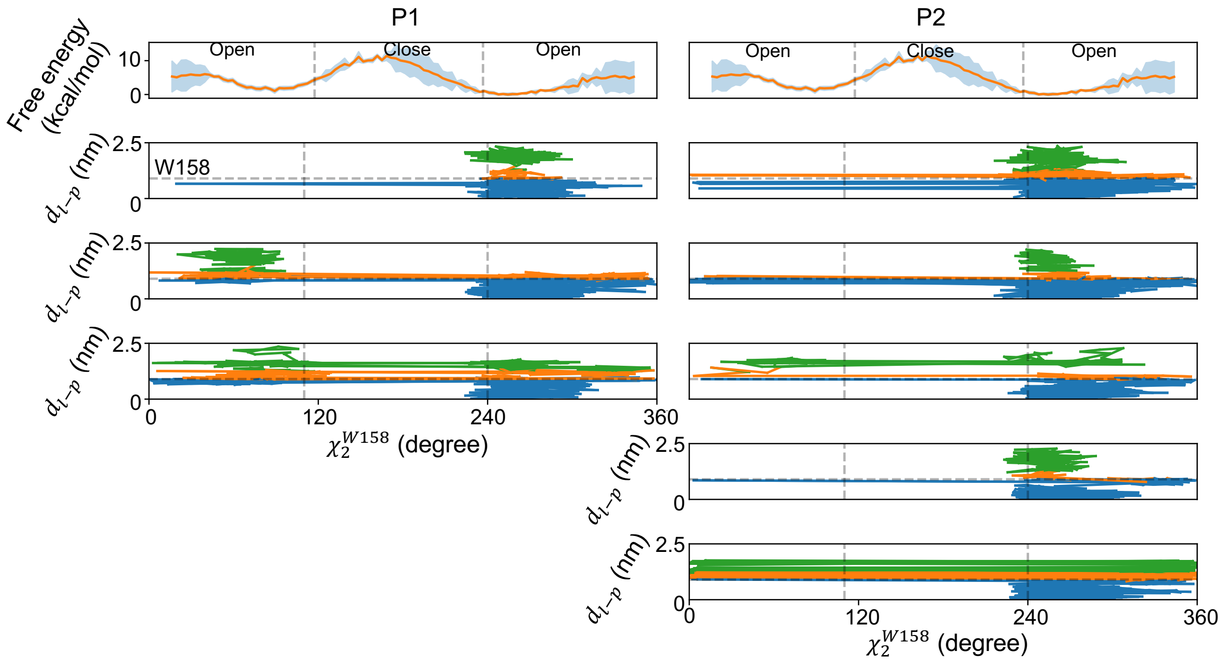


**Figure S28:** Projection of DEET dissociation trajectories onto $\chi_2^{W158}$ and $d_{l-p}$ coordinates. 8 OPES-Flooding dissociation trajectories are projected onto the $\chi_2^{W158}$ dihedral angle and the $d_{l-p}$ coordinate and compared with the corresponding $\chi_2^{W158}$ free energy landscape. Trajectory segments are colored blue when the ligand does not cross W158, orange during W158 crossing, and green after complete crossing of W158.

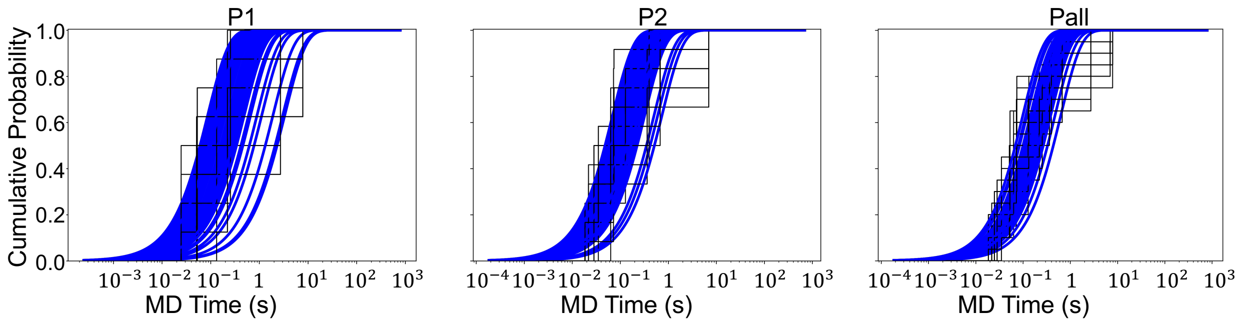


**Figure S29:** Kolmogorov-Smirnov test for dissociation pathways using 50 bootstrap samples for MhOR5-EOL.

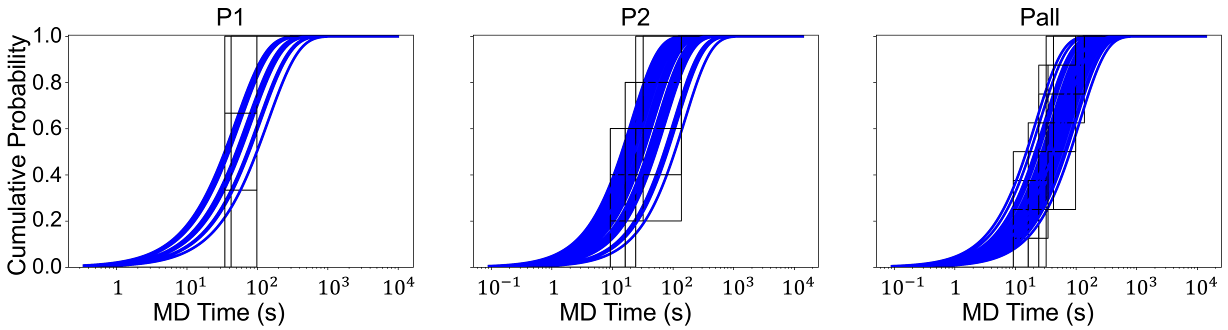


**Figure S30:** Kolmogorov-Smirnov test for dissociation pathways using 50 bootstrap samples for MhOR5-DEET.

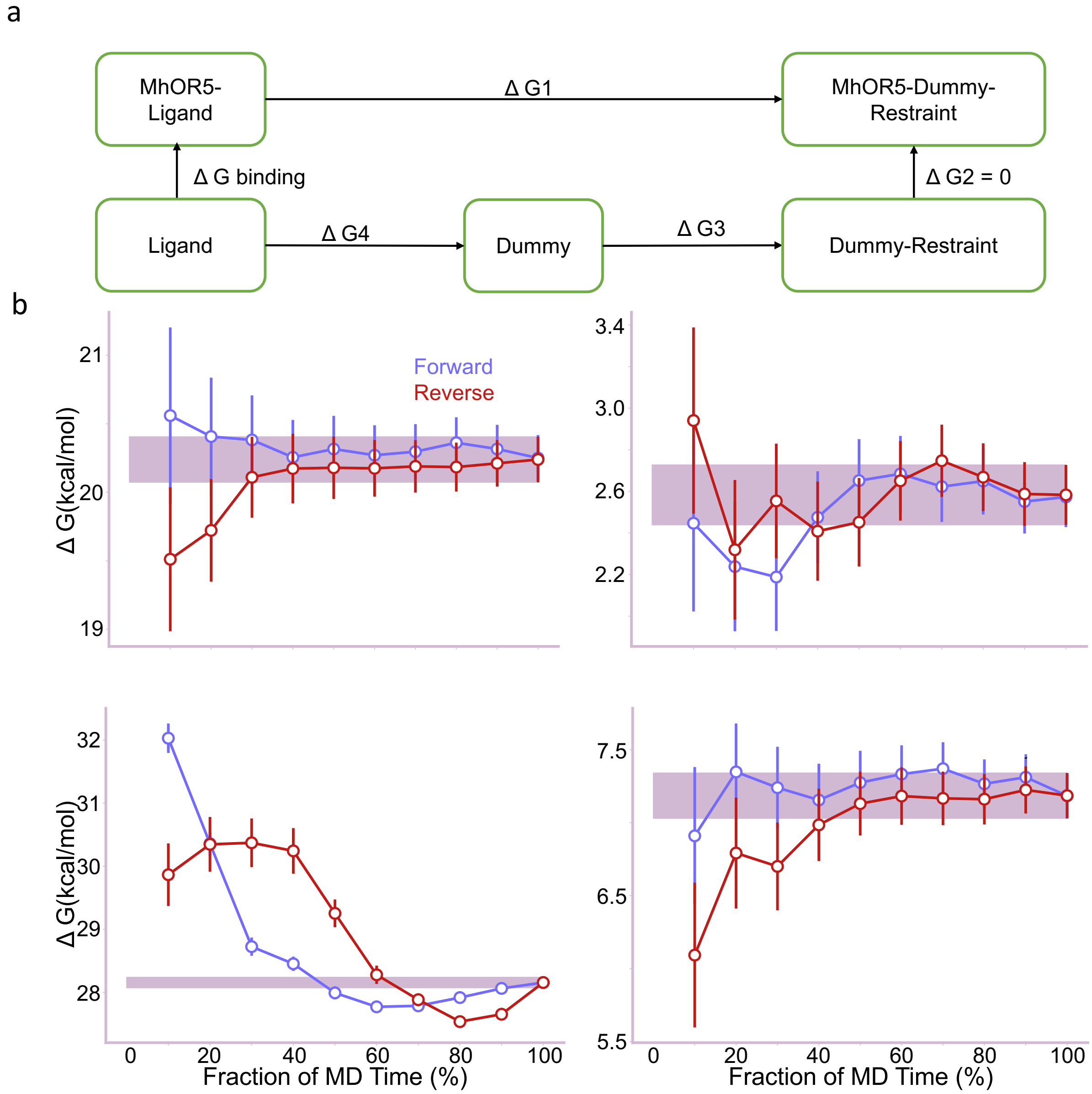


**Figure S31:** Schematic of the thermodynamic cycle and convergence assessment of FEP calculations. (a) Free energy perturbation (FEP) thermodynamic cycle used for absolute binding free energy (ABFE) calculations. (b) Time evolution of $\Delta G$ and its convergence behavior. The horizontal axis represents the fraction of the total simulation time, and the final uncertainty is indicated by the pink shaded region. Panels are shown from top left to bottom right for the EOL complex, EOL ligand-only system, DEET complex, and DEET ligand-only system.

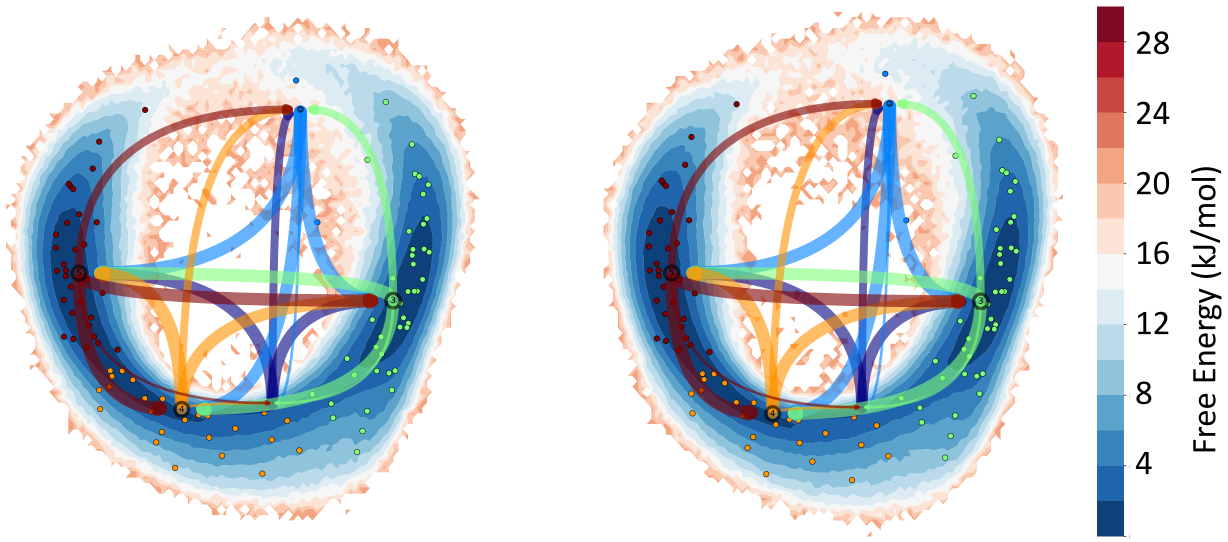


**Figure S32:** HMR Kinetic Validation. Markov state models of the alanine dipeptide show identical metastable states and transition kinetics with and without HMR at a 2 fs timestep. Projections of no HMR (left) and HMR (right) onto the first two dimensions of TIC: free energy topography, macrostate population, and transition rate. Each small circle represents a microstate. Each large circle represents a macrostate, including the microstates of the same color, and its size is proportional to the population of that state. The lines represent the transition rate between every two macrostates, and the line thickness is proportional to $\log_2(1/\mathrm{MFPT}_{ij})$.
